# Low FoxO expression in *Drosophila* somatosensory neurons protects dendrite growth under nutrient restriction

**DOI:** 10.1101/735605

**Authors:** Amy R. Poe, Yineng Xu, Christine Zhang, Kailyn Li, David Labib, Chun Han

## Abstract

During prolonged nutrient restriction, developing animals redistribute vital nutrients to favor brain growth at the expense of other organs. In *Drosophila*, such brain sparing relies on a glia-derived growth factor to sustain proliferation of neural stem cells. However, whether other aspects of neural development are also spared under nutrient restriction is unknown. Here we show that dynamically growing somatosensory neurons in the *Drosophila* peripheral nervous system exhibit organ sparing at the level of arbor growth: Under nutrient stress, sensory dendrites preferentially grow as compared to neighboring non-neural tissues, resulting in dendrite overgrowth. Underlying this neuronal nutrient-insensitivity is the lower expression of the stress sensor FoxO in neurons. Consequently, nutrient restriction suppresses Tor signaling less and does not induce autophagy in neurons. Preferential dendrite growth is functional desirable because it results in heightened animal responses to sensory stimuli, indicative of a potential survival advantage under environmental challenges.

## INTRODUCTION

Proper animal development requires coordinated growth of various organs, so that correct relative organ proportions can be reached in mature individuals. However, when developing animals face severely adverse conditions, such as limited availability of nutrients, they reallocate essential resources to favor growth of vital organs at the expense of other organs. This phenomenon of “organ sparing” is exemplified by the preferential growth of the brain in human fetuses experiencing intrauterine growth restriction, resulting in small-size newborns with relatively large heads (Gruenwald, 1963). How systemic growth control is altered at the molecular level to spare the brain remains poorly understood.

A well characterized example of brain sparing occurs in *Drosophila* larvae experiencing nutrient deprivation. Systemic larval body growth of *Drosophila* is controlled by the conserved insulin/insulin-like growth factor (IGF) pathway (Rulifson et al., 2002). *Drosophila* insulin-like peptides (Dilps) secreted by the insulin producing cells (IPCs) in the larval brain promote cell proliferation and growth of peripheral tissues by activating the insulin receptor (InR) and the downstream signaling components phosphatidylinositol 3-kinase (PI3K) and Akt (PKB) (Verdu et al., 1999; Brogiolo et al., 2001; Ikeya et al., 2002; Oldham et al., 2002). Nutrient restriction suppresses insulin secretion through an intricate nutrient sensing mechanism involving inter-organ communications between the fat body and IPCs (Ikeya et al., 2002; Geminard et al., 2009; Rajan and Perrimon, 2012), and consequently, curbs the growth of most peripheral tissues. However, the larval brain is protected against nutrient deprivation and exhibits continuous neurogenesis (Cheng et al., 2011). This protection is mediated by the glia-derived Jelly belly (Jeb) ligand that activates the Anaplastic lymphoma kinase (Alk) receptor on neural stem cells (NSCs) to turn on the downstream PI3K pathway independent of nutrition (Cheng et al., 2011). Although cell proliferation of the nervous system is spared under nutrient deprivation, whether other aspects of neural development are also subject to organ sparing is unknown.

The arbor expansion of post-mitotic neurons involves only increases of cell size and therefore represents a different type of neural growth from cell proliferation. Upon initial innervation of the target field, the dendritic or axonal arbor of the neuron expands in coordination with the tissue it innervates. For example, the dendritic arbors of *Drosophila* somatosensory neurons called dendritic arborization (da) neurons are known to scale with the body wall during normal larval development (Parrish et al., 2009). This scaling involves size increases of individual epidermal cells and simultaneous expansion of da dendritic arbors, such that neurons maintain the same coverage of the sensory fields while the body surface area expands exponentially (Jiang et al., 2014). Da neurons are categorized into four classes that differ in their dendrite morphology and transcription factor expression (Grueber et al., 2002; Hattori et al., 2013). Recently, class IV da (C4da) neurons, which completely cover the body surface and thus are called “space-filling” neurons (Grueber et al., 2002; Grueber et al., 2003), were found to elaborate more dendrite branches when larvae develop on a low-nutrient diet (Watanabe et al., 2017), suggesting that dendritic scaling of C4da neurons is regulated by the nutrient state. However, whether this dendritic hyperarborization is related to organ sparing and how nutrient stress promotes dendrite growth are unclear.

Dendrite growth in both insects and mammals is positively regulated by the conserved PI3K-Akt-mechanistic target of rapamycin (mTOR) pathway (Jaworski et al., 2005; Kumar et al., 2005; Parrish et al., 2009; Skalecka et al., 2016). Receiving signaling inputs from membrane receptor tyrosine kinases (RTKs), notably InR (Sancak et al., 2007; Vander Haar et al., 2007; Wang et al., 2007), this pathway enhances translation in most cells by mTOR kinase-mediated phosphorylation of S6 protein kinase (S6K) and 4E-binding protein (4E-BP) (Burnett et al., 1998). At the center of this pathway, mTOR activity is also influenced by the cellular state, including nutrient availability, cellular energy levels, and stress factors (Zoncu et al., 2011). In particular, cellular nutrient starvation suppresses mTOR and consequently induces autophagy (Ganley et al., 2009; Hosokawa et al., 2009; Jung et al., 2009), the self-eating process that helps to conserve and recycle vital cellular building blocks. mTOR regulates autophagy partially through the transcription factor EB (TFEB), which promotes autophagosome biogenesis but is suppressed by mTOR-mediated phosphorylation (Jung et al., 2009; Martina et al., 2012; Roczniak-Ferguson et al., 2012). Among the cellular stress sensors that inhibit mTOR activity, the forkhead box O (FoxO) family of transcription factors can by activated by a variety of stress signals and respond by suppressing cell growth and inducing autophagy (Eijkelenboom and Burgering, 2013). Although the regulation of mTOR activity by cellular stress has been extensively investigated in many cell types, how mTOR signaling is modulated by the nutrient state to impact neuronal arbor growth has not been examined. Furthermore, although FoxO members have been found to enhance dendritic space-filling of C4da neurons in *Drosophila* (Sears and Broihier, 2016) and to regulate dendrite branching and spine morphology of adult-generated neurons in mice (Schaffner et al., 2018), whether they also influence neuronal arbor growth in response to nutrient stress is unclear.

In this study, we demonstrate that dynamically growing *Drosophila* da neurons exhibit organ sparing at the level of individual cells; that is, dendrites grow preferentially at the expense of other non-neural tissues under nutrient stress. Mechanistically, the amplitude of Tor signaling in da neurons is much less affected by nutrient stress than in non-neural tissues like epidermal cells. As a result, under these conditions da neurons do not undergo autophagy as do surrounding tissues, and the neurons display a relative growth advantage. Underlying the distinct sensitivities to nutrient stress are the differential FoxO expression levels in da neurons and epidermal cells: Foxo is expressed at too low levels in neurons to significantly affect dendrite growth, while it is expressed at a much higher level in epidermal cells, resulting in suppression of cell growth only under nutrient restriction. Functionally, preferential dendrite growth of da neurons increases the sensory acuity, allowing larvae to respond more nimbly to environmental stimuli.

## RESULTS

### Nutrient restriction affects the growth of epidermal cells and C4da neurons differentially

A recent study by Watanabe et al. (2017) reported that C4da neurons were hyperarborized when *Drosophila* larvae developed on a low yeast diet that restricts the availability of lipids and amino acids, an interesting and surprising finding that agrees with our independent observation. In our experiments, we examined larvae reared in high yeast (HY, 8% yeast) and low yeast (LY, 1% yeast) media that otherwise contained only glucose as a carbon source. In late third instar larvae, C4da neurons in the HY condition showed sparse dendrites with gaps of dendritic coverage between neighboring neurons (Figure 1A). In contrast, C4da neurons in the LY condition completely covered the body wall with dense dendrites (Figure 1B).

**Figure 1.**
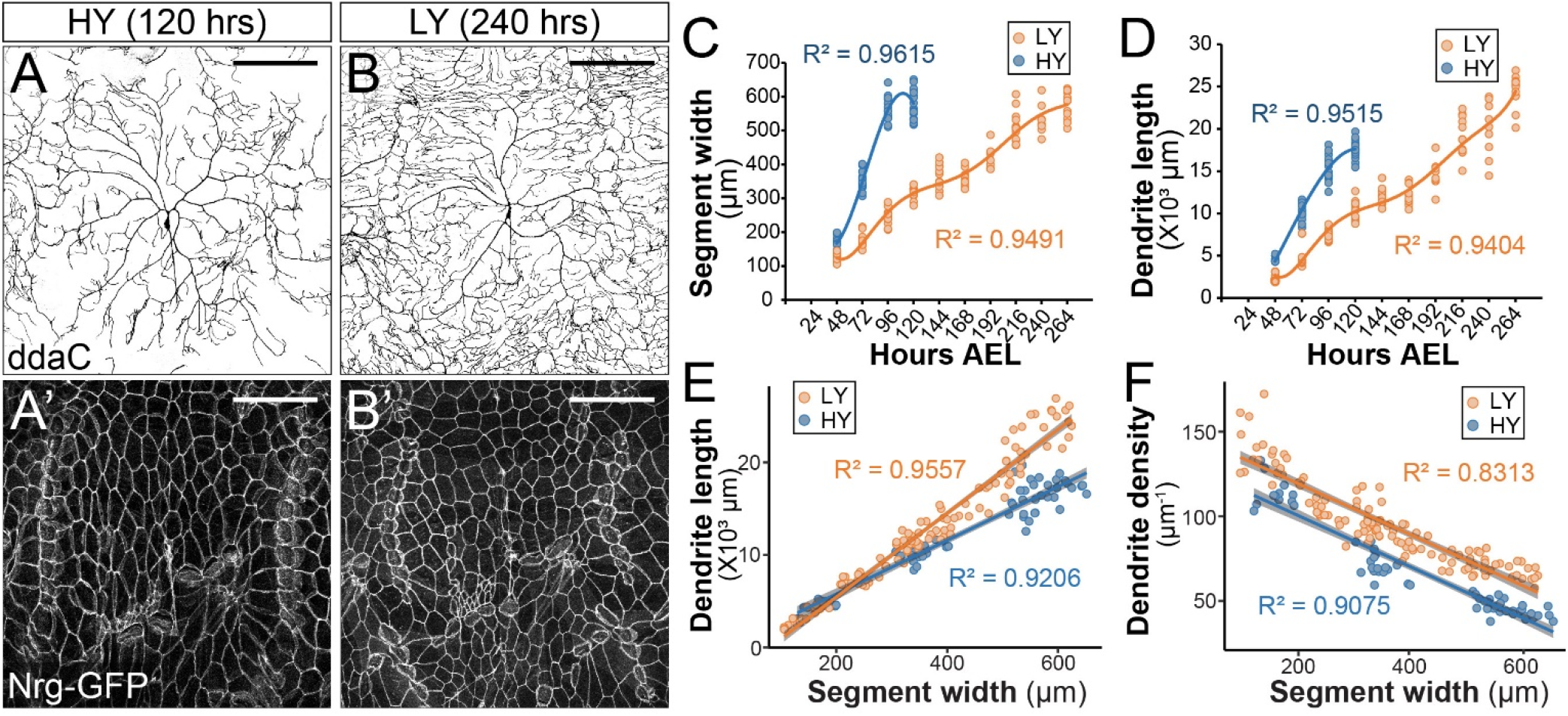
Nutrient restriction affects the growth of epidermal cells and C4da neurons differentially. (A-B’) Double labeling of ddaC neurons by *ppk-CD4-tdTom* (A and B) and epidermal cells by the septate junction marker Nrg-GFP (A’ and B’) in the high yeast (HY, 8%) condition at 120 hrs after egg laying (AEL) (A and A’) and in the low yeast (LY, 1%) condition at 240 hrs AEL (B and B’). (C and D) Plots of segment width (C) and total dendrite length of ddaC neurons (D) versus time in HY and LY conditions. (E and F) Plots of total dendrite length (*p*≪0.05) (E) and dendrite density (total dendrite length/dendrite coverage area, *p*≪0.05) (F) with segment width in HY and LY conditions. Each circle represents a segment in (C) and a ddaC neuron in (D-F); n=63 for HY; n= 115 for LY. Solid lines represent polynomial fits in (C) and (D) and linear fits in (E) and (F). R^2^ represents coefficient of determination of the linear regression. Grey shading in (E) and (F) represents a 0.95 confidence interval (CI) of the linear model. *p*-value represents the possibility that the slops of two yeast conditions are the same. Scale bars, 100 μm.

To understand the molecular and cellular basis of this nutrient-regulated dendrite growth, we first investigated how C4da neurons and the body wall grow differently during larval development in HY and LY media. We thus measured various parameters of the larval body wall and dendrite growth every 24 hours (hrs) starting from 48 hrs after egg laying (AEL) to the wandering 3rd instar stage (Figures 1 and S1). The body wall was measured at the levels of the whole body (body length), the body segments (segment width), and individual epidermal cells (average cell width as visualized by the septate junction marker Nrg-GFP). We found as expected that all three parameters correlated with each other in both HY and LY conditions (Figure S1G and S1H). We have therefore taken the segment width as an indicator of the larval body size.

**Figure S1.**
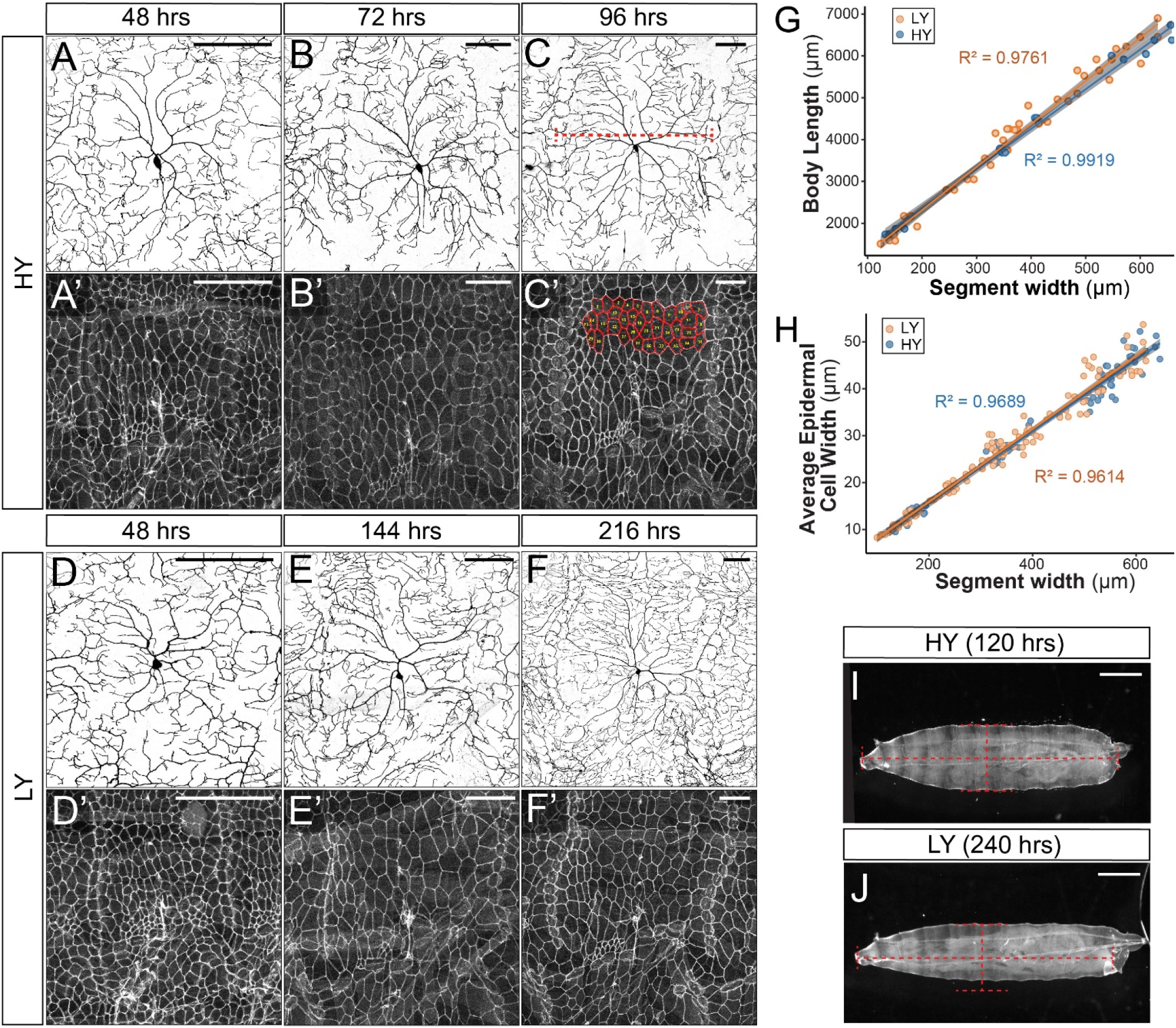
Development of C4da neurons and epidermal cells in high and low yeast media. (A-F’) Double labeling of ddaC neurons by *ppk-CD4-tdTom* (A-F) and epidermal cells by Nrg-GFP (A’-F’) in HY and LY conditions at times indicated. The red dotted line in (C) indicates how the segment width is measured. The measured epidermal cells are outlined in (C’). (G and H) Plots of body length (*p*>0.05) (G) and average epidermal cell width (p>0.05) (H) with segment width in HY and LY conditions. Each circle represents a segment; n=63 for HY; n=115 for LY. Solid lines represent linear fits. R^2^ represents coefficient of determination of the linear regression. Grey shading in (E) and (F) represents a 0.95 confidence interval (CI) of the linear model. *p*-value represents the possibility that the slops of two yeast conditions are the same. (I and J) Representative animals at 120h AEL in HY condition (I) and at 240h AEL in LY condition (J). The red dotted lines match the body width and length of the larva in HY. Scale bars, 100 μm in (A-F’), 300 μm in (I) and (J).

Compared to the HY condition, LY caused a significant delay in larval body growth, with animals in LY taking 2.2 times longer (264 hrs compared to 120 hrs) to reach a nearly identical maximum segment width of ~600 μm before pupariation (Figures 1A-1C). These observations verify that the LY medium caused the larvae to experience nutrient stress and developmental delay. The fact that the pupariation occurs at the same maximum segment width regardless of the rate of growth indicates that the segment width is not only a good proxy for body size, but it also provides a good indication of the development stage.

C4da neurons also grew slower in LY than in HY, as indicated by smaller increases of the total dendrite length during each 24 hr period (Figure 1D). However, because LY larvae had more time to develop, their neurons ultimately outgrew those of HY larvae, resulting in 1.39-fold longer total dendrites at the end of the larval period (Figure 1D). To make more meaningful comparison of dendrite growth in animals of similar developmental stages prior to pupariation, we plotted the dendrite length against the segment width (Figure 1E), as the segment width better indicates of the developmental stage than the age of the larva. This plot shows that C4da dendrites grow 57% faster (based on the slopes of the linear fits) in LY than in HY when normalized by the segment width. Interestingly, the dendrite density (dendrite length/dendrite field size) was always higher in LY when plotted against the segment width, even though the dendrite length was not greater in LY when the segment width was below 200 μm (Figure 1F). Likely contributing to this discrepancy between dendrite density and dendrite length is that larvae reared in LY were thinner than those in HY (Figures S1I and S1J) and consequently had smaller body wall areas to cover by C4da dendrites.

Collectively, these results show that while nutrient restriction delays the overall larval growth, it slows down the growth of C4da neurons and non-neural tissues to different degrees, such that C4da neurons exhibit a growth advantage, or are spared, under nutrient stress.

### The InR-Tor pathway mediates the preferential dendrite growth under nutrient stress

InR and Tor (the *Drosophila* mTOR) are the main players through which nutrient availability controls larval systemic growth (Boulan et al., 2015). To compare dendrite growth in larvae reared in HY and LY media, we measured the normalized dendrite length (dendrite length/segment width) when larvae reached a segment width between 500 and 550 μm. Downregulation of the InR/Tor pathway in neurons by *InR* knockdown (InR RNAi), *InR^DN^* (dominant negative) overexpression (*InR* DN), *Tor* knockdown (Tor RNAi), or *Tor^DN^* overexpression (Tor DN) in the HY condition caused mild or statistically insignificant dendrite reduction compared to the control (Figures 2A-C, 2G, and S2A-S2C). In contrast, the same genetic manipulations caused pronounced dendritic reduction in the LY condition, generating dendrite patterns resembling those of wildtype neurons in the HY condition (Figures 2D-F, 2G, and S2D-S2F). The ratio of the average normalized dendrite length between LY and HY thus drops from 1.44 for control neurons to values closer to 1 for neurons in which *InR* or *Tor* is suppressed (Figure 2H). These results suggest that neuronal InR/Tor signaling is responsible for the preferential dendrite growth observed under nutrient stress.

**Figure 2.**
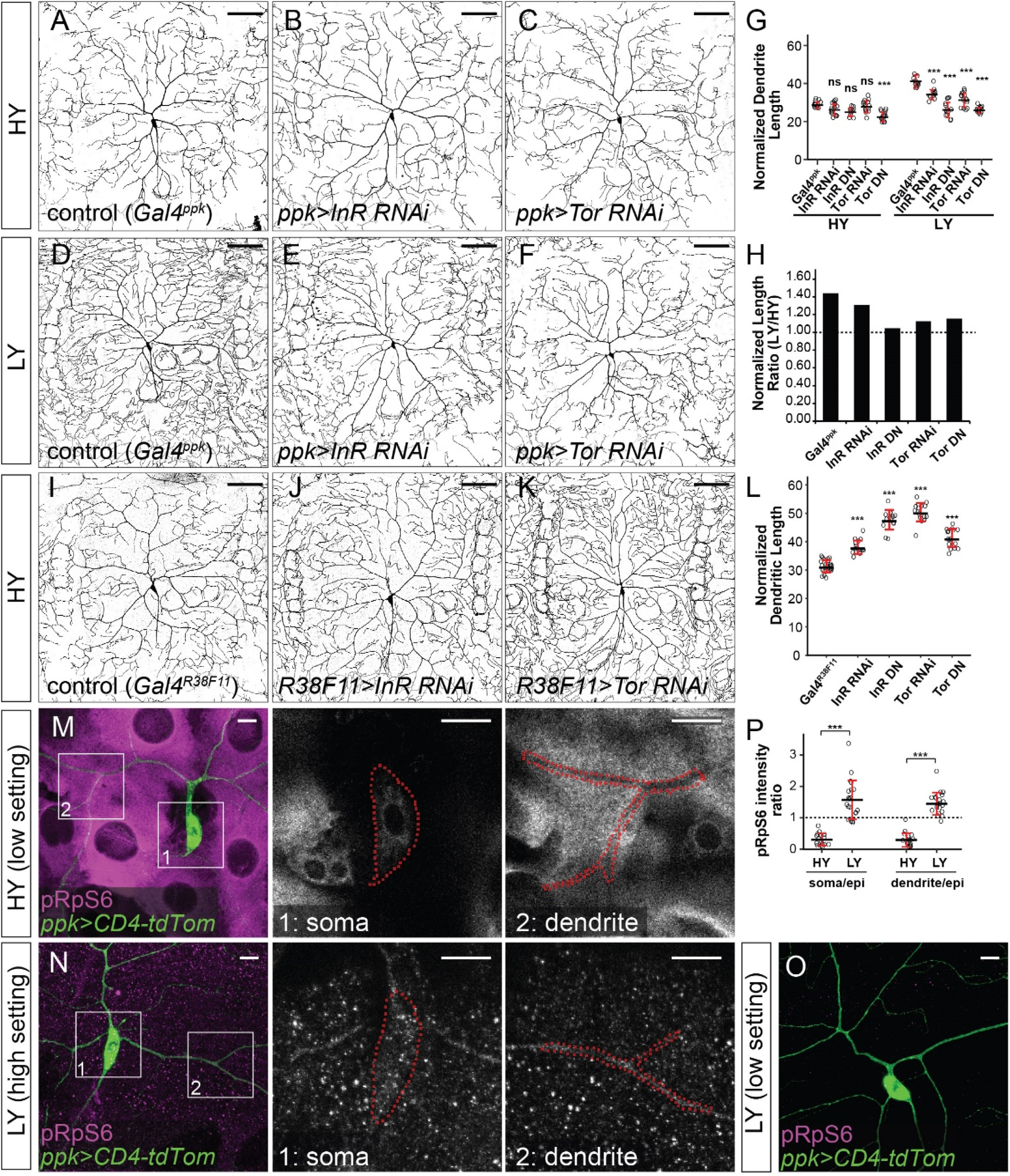
The InR-Tor pathway mediates the preferential dendrite growth under nutrient stress. (A-F) ddaC neurons in the *Gal4^ppk^* control (A and D) and animals expressing *Gal4^ppk^*-driven *InR* RNAi (B and E) and *Tor* RNAi (C and F) in HY and LY conditions. (G) Quantification of normalized dendrite length (total dendrite length/segment width) in HY and LY conditions. HY: n=14 for *Gal4^ppk^*, n=15 for *InR* RNAi, n=11 for *InR* DN, n=15 for *Tor* RNAi, n=15 for *Tor* DN; LY: n=14 for *Gal4^ppk^*, n=12 for *InR* RNAi, n=14 for *InR* DN, n=15 for *Tor* RNAi, n=15 for *Tor* DN. (H) The ratios of average normalized dendrite length between LY and HY. (I-K) ddaC neurons in the *Gal4^R38F11^* control (I) and animals expressing *Gal4^R38F11^*-driven *InR* RNAi (J) and *Tor* RNAi (K) in HY condition. (L) Quantification of normalized dendrite length in HY condition. n=23 for *Gal4^R38F11^*, n=17 for *InR* RNAi, n=17 for *InR* DN, n=16 for *Tor* RNAi, n=17 for *Tor* DN. (M-O) pRpS6 staining (magenta) of ddaC neurons (Green) and epidermal cells in HY and LY conditions in 2-dimesional (2D) projections. The insets in (M) and (N) show pRpS6 staining at the soma (1) and primary dendrites (2) in single confocal sections, with the somas and dendrites outlined. High settings and low settings stand for high and low pRpS6 detection settings. (P) Quantification of pRpS6 intensity ratio (soma/epidermal cells and dendrites/epidermal cells) in HY and LY conditions. Soma/epi: n=17 for HY, n=20 for LY; dendrites/epi: n=16 for HY, n=20 for LY. For all quantifications, \*\*\**p*<0.001; ns, not significant; One-way ANOVA and Tukey’s HSD test; each circle represents a neuron. Black bars, mean; red bars, SD. Scale bars, 100 μm in (A-K); 10 μm in (M-O).

**Figure S2.**
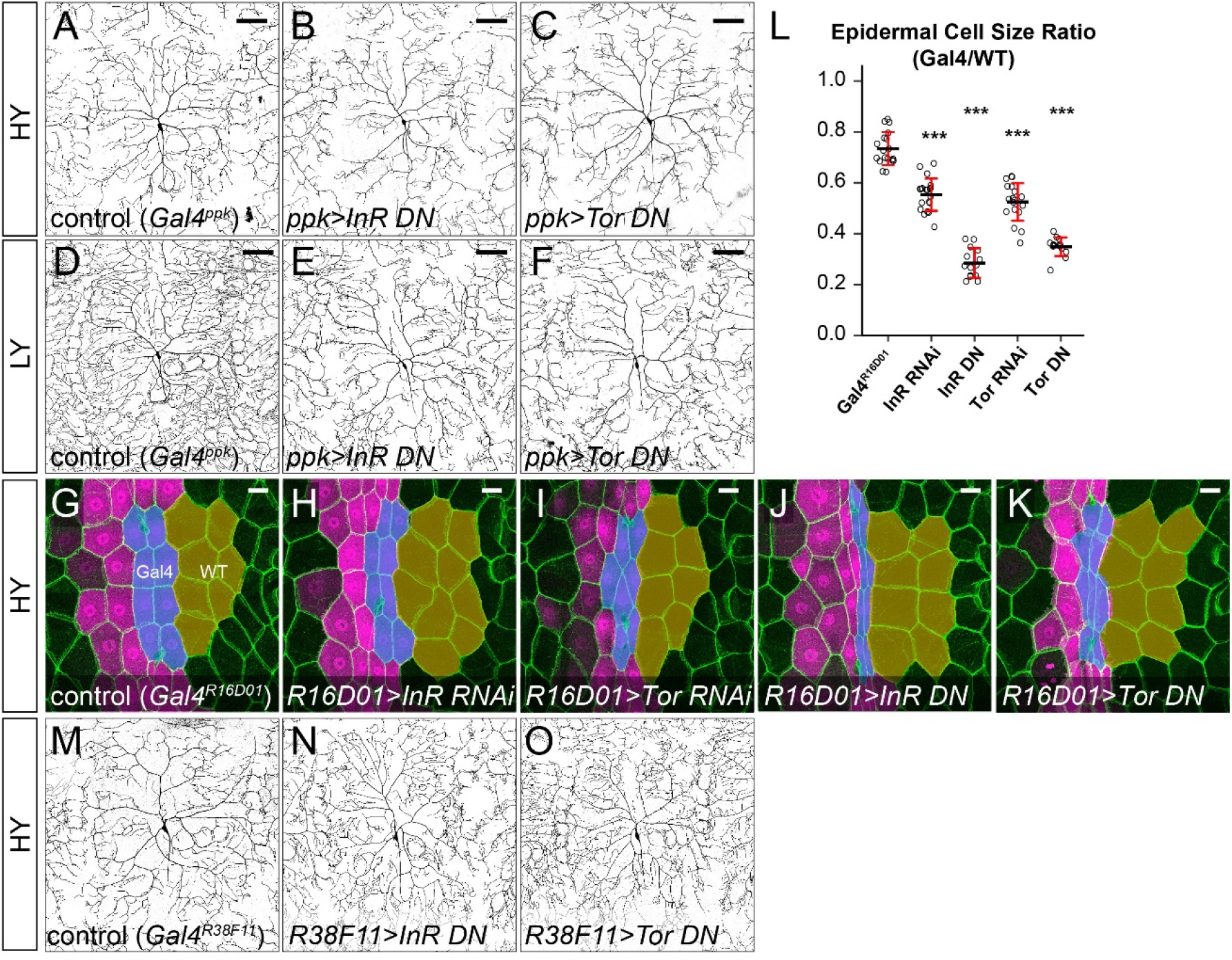
The effects of suppressing *InR* and *Tor* in ddaC neurons and epidermal cells in HY and LY conditions. (A-F) ddaC neurons in the *Gal4^ppk^* control (A and D) and animals expressing *Gal4^ppk^*-driven *InR* DN (B and E) and *Tor* DN (C and F) in HY and LY conditions. (G-K) Epidermal cells in the *Gal4^R16D01^* control (G) and animals expressing *Gal4^R16D01^*-driven *InR* RNAi (H), *Tor* RNAi (I), *InR* DN (J) and *Tor* DN (K) in the HY condition. *Gal4^R16D01^* domain is labeled by mIFP expression (magenta). All epidermal cells are labeled by Nrg-GFP (green). The blue and yellow overlays indicate the measured Gal4-expressing and wildtype (WT) epidermal cells, respectively. (L) Quantification of epidermal cell size ratio (*Gal4^R16D01^* cells/WT cells). Each circle represents a segment; n=18 for *Gal^R16D01^*, n=20 for *InR* RNAi, n=14 for *InR* DN, n=18 for *Tor* RNAi, n=14 for *Tor* DN. \*\*\**p*<0.001; ns, not significant; One-way ANOVA and Tukey’s HSD test. Black bars, mean; red bars, SD. (M-O) ddaC neurons in the *Gal4^R38F11^* control (M) and animals expressing *Gal4^R38F11^*-driven *InR* DN (N) and *Tor* DN (O) in the HY condition. (A), (D), (M) are the same as Figure 2A, 2D, and 2I, respectively. Scale bars, 100 μm in (A-F), 50 μm in (M-O) and (G-K).

To test whether slowing down epidermal growth, such as seen under nutrient stress, can lead to excessive dendrite growth, we inhibited *InR* and *Tor* in epidermal cells under the HY condition. *UAS*-driven transgenes were expressed by *Gal4^R16D01^* in a stripe of epidermal cells in the middle of each segment (Poe et al., 2017), so that the Gal4-expressing epidermal cells can be compared to neighboring wildtype cells. Downregulating *InR* or *Tor* effectively reduced the epidermal cell size as expected (Figure S2G-S2K), reducing the ratio between the sizes of Gal4-positive (Gal4) and neighboring Gal4-negative (WT) cells (Figure S2L). When *InR* or *Tor* was suppressed in the whole epidermal sheet by using the pan-epidermal driver *Gal4^R38F11^*, the larvae took 6-30 extra hours to reach the segment width of 500-550 μm and exhibited 22%-61% greater normalized dendrite length than controls (Figures 2I-2L and S2M-S2O). Taken together, above results in Figure 2 and Figure S2 demonstrate that changes in the relative strengths of the InR/Tor signaling in neurons and epidermal cells can alter dendritic scaling: Reducing the throughput of the pathway in neurons can cause dendrite reduction, while suppression of InR/Tor in the epidermis can lead to dendrite overgrowth.

To determine how the nutrient level modulates InR/Tor signaling in neurons and epidermal cells, we examined Tor activity by immunostaining phosphorylated Ribosomal protein S6 (pRpS6), a substrate of S6K and a faithful indicator of Tor activity in *Drosophila* tissues (Ruvinsky and Meyuhas, 2006; Kim et al., 2017). In HY, the cytoplasm of epidermal cells exhibited high and uniformly distributed pRpS6 signals, while the soma and primary dendrites of C4da neurons showed comparatively low pRpS6 intensities (Figure 2M). As a result, the ratios of pRpS6 intensity (average intensity within regions of interest) between the neuronal compartments and epidermal cells are much lower than 1 (Figure 2P). Under nutrient stress, the overall pRpS6 staining on the larval body wall dramatically decreased (Figure 2O), consistent with the notion that nutrient stress reduces InR/Tor signaling in peripheral tissues (Geminard et al., 2009). However, in these animals, the pRpS6 signals were brighter and more even in C4da cell bodies and dendrites than in the epidermal cells (Figure 2N), causing the ratios of pRpS6 intensity between the neuronal compartments and epidermal cells to be larger than 1 (Figure 2P). These data suggest that nutrient stress switches the relative strength of the InR/Tor signaling in C4da neurons and epidermal cells such that the neurons gain a growth advantage over epidermal cells.

### The lack of autophagy induction protects C4da neuronal growth under nutrient stress

Nutrient stress suppresses mTOR activity to induce many cellular responses, including autophagy (He and Klionsky, 2009). We therefore tested if autophagy is also differentially regulated by the nutrient level in C4da neurons and epidermal cells. To examine the autophagy level, we used an mCherry-Atg8a reporter under the control of the endogenous Atg8a regulatory sequence, which labels autophagic structures (Hegedus et al., 2016). As expected, the autophagosome level in epidermal cells was low under the HY condition (Figure 3A) but increased 5 folds in the LY diet (Figures 3B and 3C). In contrast, autophagosomes in C4da cell bodies kept at low levels under both HY and LY conditions (Figures 3A-3C). Similarly, Lamp-mCherry, a lysosomal and autolysosomal marker driven by the endogenous *Lamp1* promoter (Hegedus et al., 2016), showed a 4.8-fold increase of signals in epidermal cells under nutrient stress (Figures S3A-S3C). In contrast, the same reporter exhibited a much lower baseline labeling in C4da cell bodies (12.6% of that in epidermal cells) under HY and a milder (2.74-fold) increase under LY (Figures S3A-S3C). To examine the levels of autophagic flux, we overexpressed in epidermal cells and neurons a tandem fluorescent marker GFP-mCherry-Atg8a, which is converted from dual colors to mCherry alone when autophagosomes mature (Kimura et al., 2007; Nezis et al., 2010). Thus, increased autophagic flux results in a reduction of cytosolic GFP and an increase of vesicular mCherry, indicated by an increased ratio of mCherry area over GFP intensity. Using this marker, we found that the autophagic flux level was always higher in epidermal cells and that nutrient stress enhanced autophagic flux in both epidermal cells and C4da neuron cell bodies (Figures S3D-S3I). These data suggest that C4da neurons keep the autophagy level low even under nutrient stress, despite an increase of autophagic flux.

**Figure 3.**
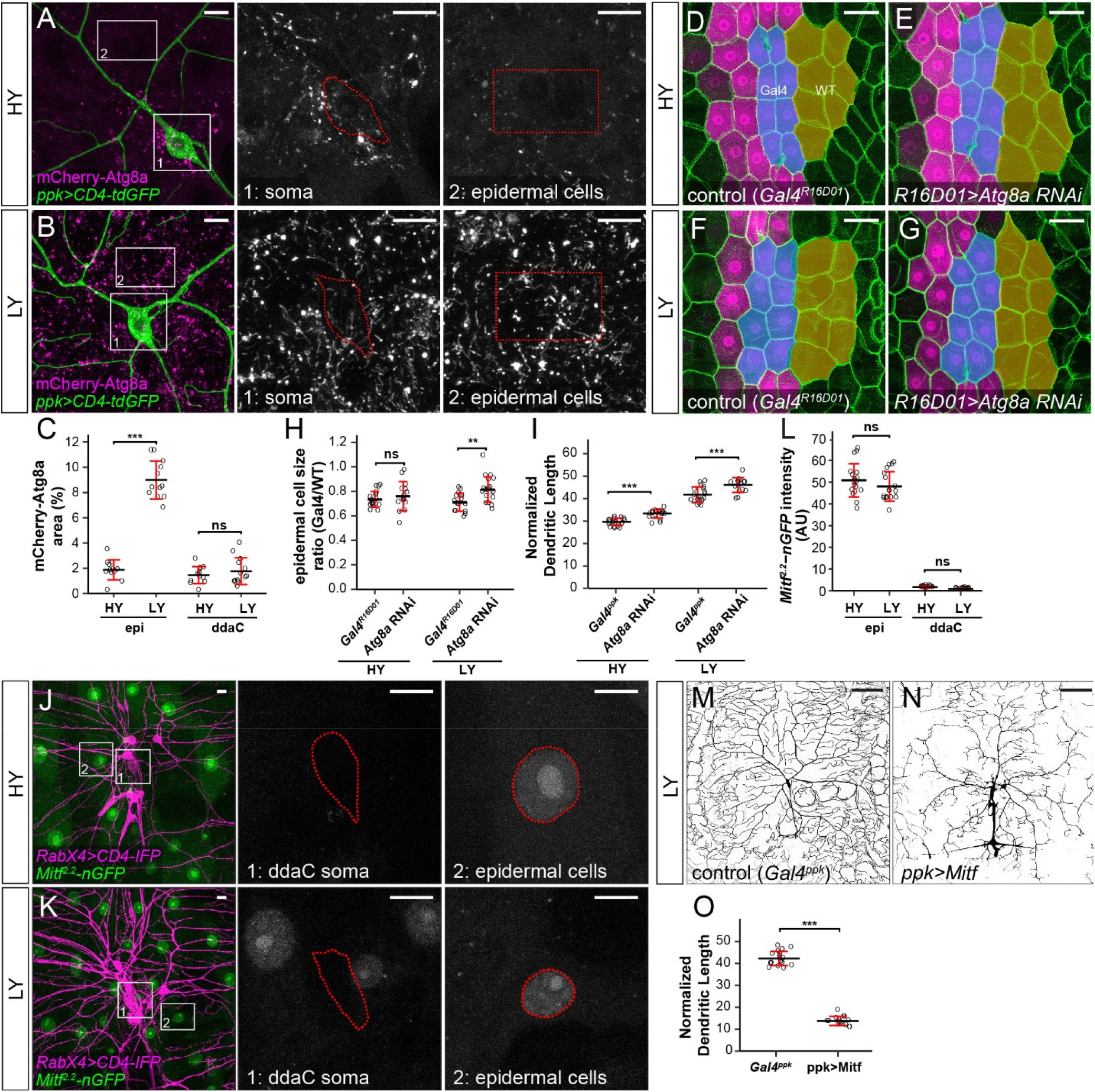
The lack of autophagy induction protects C4da neuronal growth under nutrient stress. (A and B) mCherry-Atg8a (magenta) in ddaC soma (green) and epidermal cells in HY and LY conditions. The insets in (A) and (B) show mCherry-Atg8a at the soma of ddaC (1) and epidermal cells (2). The dotted lines indicate the somas (1) and measured epidermal regions (2). All images are 2D projections. (C) Quantification of mCherry-Atg8a levels in ddaC soma and epidermal cells in HY and LY conditions, measured by the area percentage of mCherry-Atg8a-positive vesicles. Epi: n=12 for HY, n=13 for LY; ddaC: n=12 for HY, n=13 for LY. (D-G) Epidermal cells in the *Gal4^R16D01^* control and animals expressing *Gal4^R16D01^*-driven *Atg8a* RNAi in HY and LY conditions. *Gal4^R16D01^* domain is labeled by mIFP expression (magenta). All epidermal cells are labeled by Nrg-GFP (green). The blue and yellow overlays indicate the measured Gal4-expressing and wildtype (WT) epidermal cells, respectively. (D) is the same as Figure S2G. (H) Quantification of epidermal cell size ratio (*Gal4^R16D01^* cells/WT cells) in HY and LY conditions. Each circle represents a segment; HY: n=18 for control, n=14 for Atg8a RNAi; LY: n=16 for control, n=19 for Atg8a RNAi. (I) Quantification of normalized dendritic length in control and *Atg8a* RNAi animals in HY and LY conditions. Each circle represents a neuron; HY: n=17 for *Gal4^ppk^*, n=16 for *Atg8a* RNAi; LY: n=15 for *Gal4^ppk^*, n=12 for *Atg8a* RNAi. *Gal4^ppk^* is the same dataset as figure 2G (J and K) *Mitf^2.2^-nGFP* (green) in ddaC neurons (magenta) and epidermal cells in HY and LY conditions. The insets in (A) and (B) show *Mitf^2.2^-nGFP* at ddaC somas (1) and epidermal cells (2), with the somas and epidermal cell nuclei outlined. All images are 2D projections. (L) Quantification of *Mitf^2.2^-nGFP* intensity in ddaC neuron soma and epidermal cells in HY and LY conditions. epi: n=18 for HY, n=17 for LY; ddaC: n=17 for HY, n=17 for LY. (M and N) DdaC neurons in the *Gal4^ppk^* control (M) and animals expressing *Gal4^ppk^*-driven *Mitf* (N) in LY condition. (M) is the same as Figure 2D. (O) Quantification of normalized dendritic length in control and *Mitf* overexpression animals in LY condition. Each circle represents a neuron; n=15 for *Gal4^ppk^* 1, n=18 for *ppk*> *Mitf. Gal4^ppk^* is the same dataset as figure 2G. For all quantifications, \*\*\**p*<0.001; \*\**p*<0.01; ns, not significant; One-way ANOVA and Tukey’s HSD test. Black bars, mean; red bars, SD. Scale bars, 10 μm in (A), (B), (J), and (K); 50 μm in (D-G); 100 μm in (M) and (N).

To understand the effects of autophagy on the growth of epidermal cells and neurons, we knocked down *Atg8a* to suppress autophagy. Epidermal *Atg8a* knockdown using *Gal4^R16D01^* did not have an effect on the cell size under the HY condition (Figures 3D, 3E, and 3H), consistent with the low autophagy level in these cells. However, the same manipulation caused 16% increase of the epidermal cell size under LY (Figures 3F, 3G, & 3H), supporting that idea that autophagy suppresses cell growth under nutrient stress. In comparison, neuronal *Atg8a* knockdown in both HY and LY conditions resulted in similar increases of normalized dendrite length (13% and 11% increases, respectively) (Figures S3J-S3M and 3I), suggesting that C4da neurons maintain a constant level of autophagy regardless of nutrient availability to mildly suppresses dendrite growth.

Nutrient restriction upregulates expression of autophagy-related genes through the transcription factor TFEB, a substrate of the Tor kinase (Fullgrabe et al., 2016). To understand why nutrient stress fails to induce autophagy in neurons, we examine the expression of *Mitf-GFPnls*, a transcription reporter for the *Drosophila TFEB* homolog *Mitf* (Zhang et al., 2015). *Mitf-GFPnls* showed nutrient-independent expression in epidermal nuclei but its expression could not be detected in da neurons in either HY or LY conditions (Figures 3J-3L), suggesting that da neurons possibly lack this essential transcriptional inducer of autophagy. We next investigated the effects of inducing autophagy on dendrite growth by overexpressing Mitf in C4da neurons, as overexpression of TFEB/Mitf is sufficient to dominantly induce autophagy in cells (Settembre et al., 2011; Zhang et al., 2015). This manipulation strongly reduced the normalized dendrite length under LY (Figures 3M-3O), suggesting that high autophagy levels suppress dendrite growth.

These data together suggest that nutrient restriction upregulates autophagy in epidermal cells but not in C4da neurons, and that the lack of autophagy induction likely protects neurons from growth suppression under nutrient stress.

### FoxO is differentially expressed in C4da neurons and epidermal cells to regulate their distinct responses to nutrient stress

**Figure S3.**
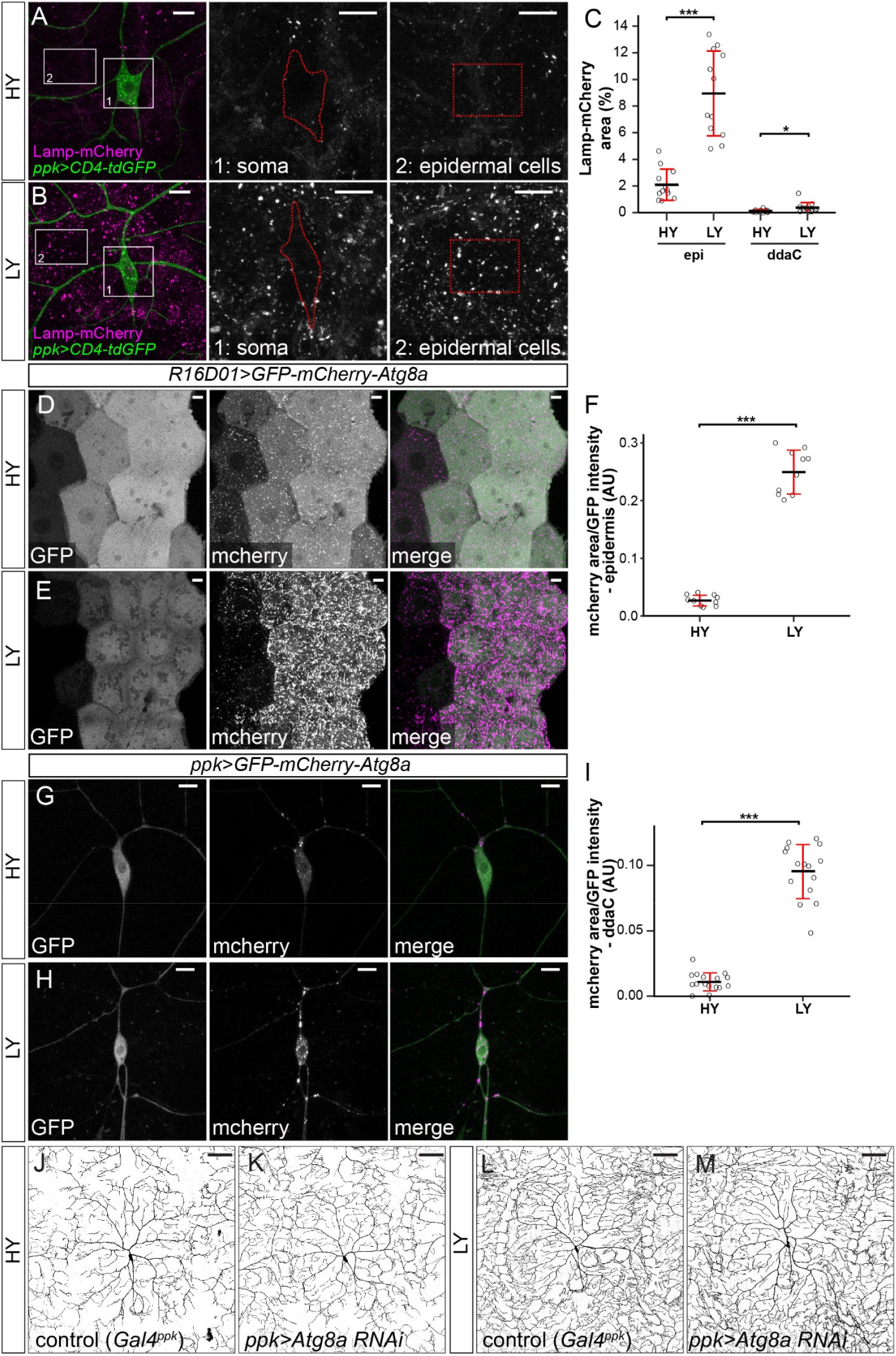
Autophagy levels in ddaC neurons and epidermal cells in HY and LY conditions. (A and B) Lamp-mCherry (magenta) in ddaC soma (green) and epidermal cells in HY and LY conditions. The insets in (A) and (B) show Lamp-mCherry at the soma of ddaC (1) and epidermal cells (2). All images are 2D projections. (C) Quantification of mChery-Atg8a levels in ddaC soma and epidermal cells in HY and LY conditions, measured by the the area percentage of Lamp-mCherry-positive vesicles. Each circle represents a segment. Epi: n=11 for HY, n=10 for LY; ddaC: n=11 for HY, n=10 for LY. (D and E) *Gal4^R16D01^*-driven *GFP-mcherry-Atg8a* in epidermal cells in HY and LY conditions. (F) Quantification of autophagic flux (area of mCherry-Atg8a-positive vesicles/GFP intensity) in epidermal cells in HY and LY conditions. Each circle represents a segment; n=10 for HY, n=10 for LY. (G and H) *Gal4^ppk^*-driven *GFP-mcherry-Atg8a* in ddaC neurons in HY and LY conditions. (I) Quantification of autophagic flux in ddaC neurons. Each circle represents a neuron; n=16 for HY, n=15 for LY. (J-M) ddaC neurons in the *Gal4^ppk^* control (J and L) and animals expressing *Gal4^ppk^*-driven *Atg8a* RNAi (K and M) in HY and LY conditions. (J) and (K) are the same as Figure 2A and 2D, respectively. For all quantifications, ***p<0.001; *p<0.05; One-way ANOVA and Tukey’s HSD test. Black bars, mean; red bars, SD. Scale bars, 10 μm in (A), (B), (D), (E), (G) and (H); 100 μm in (J-M).

To further understand why epidermal cells show stronger growth suppression than C4da neurons under nutrient stress, we tested in epidermal cells genes that potentially inhibit mTor signaling under stress conditions, including *cryptocephal* (*crc*)/*ATF4, Sestrin* (*Sesn*), and *foxo* (Eijkelenboom and Burgering, 2013; Lee et al., 2016). Expecting that inhibiting the responsible genes in epidermal cells should relieve growth suppression under nutrient restriction, we knocked down the candidate genes in epidermal cells using *Gal4^R16D01^*. Knockdown of *crc* or *Sesn* in epidermal cells did not lead to statistically significant increase of the cell size under either HY or LY conditions (Figures S4A), indicating that either these genes do not suppress epidermal cell growth or the RNAi lines were not effective. We chose to focus on *foxo* because its knockdown yielded positive results (see below).

We investigated the role of FoxO by first examining its expression in neurons and epidermal cells. A transgenic *foxo-GFP* line carries a 77 kb genomic fragment containing the full *foxo* locus with a C-terminal GFP tag and therefore should mimic the endogenous FoxO expression. FoxO-GFP showed cytoplasmic distribution in epidermal cells in both HY and LY conditions (Figures 4A-4C) but translocated to epidermal nuclei within several minutes of hypoxia (Figures S4B and S4C). In contrast, FoxO-GFP could not be detected above the background noise level in C4da cell bodies (Figures 4A-4C). To validate this tissue-specific FoxO expression pattern, we converted a MiMIC insertion line of *foxo* into a *foxo-Gal4* using the Trojan exon technique (Diao et al., 2015), so that the Gal4 is under the same transcription regulation as the endogenous *foxo* (Figure S4D). This *foxo-Gal4* drove uniform epidermal expression of a *UAS-tdTom* reporter under HY, and the expression was enhanced 2.9 folds by nutrient stress (Figures 4D-4F). In constrast, *foxo-Gal4* showed consistently minimal activities in C4da neurons under both HY and LY conditions (Figures 4D-4F). A previous study (Sears and Broihier, 2016) detected FoxO protein in C4da nuclei using antibody staining, so it is possible that FoxO is expressed at levels too low for detection in our approaches. These data suggest that FoxO is expressed in non-neural tissues like the epidermis but is absent or expressed at very low levels in C4da da neurons. We also found that *foxo-Gal4* is expressed in a subset of neurons and non-neural cells in the larval brain and ventral nerve cord (Figures S4E and S4F).

**Figure 4.**
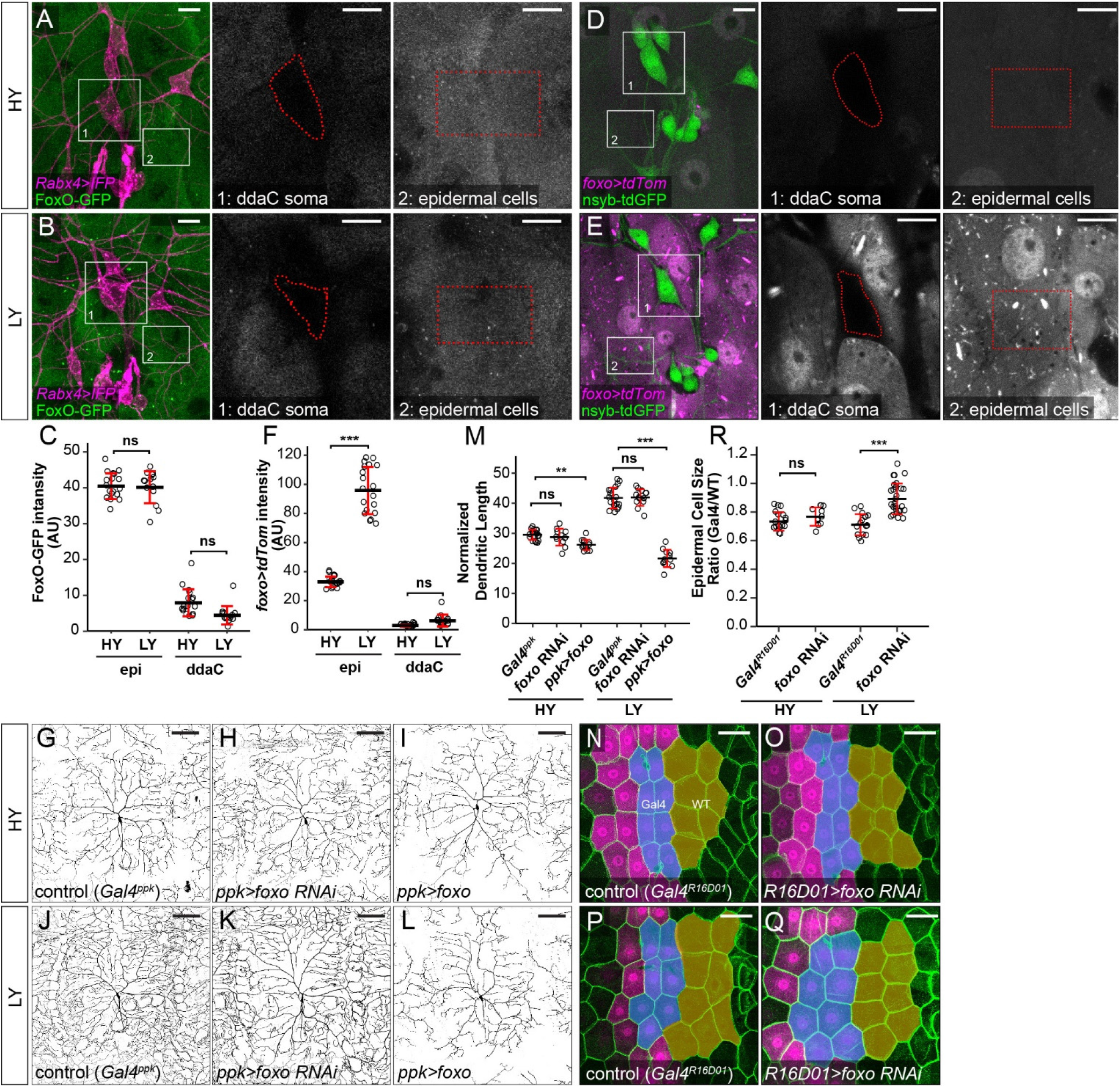
The lack of neuronal FoxO expression ensures preferential dendrite growth under nutrient stress. (A and B) FoxO-GFP (green) in da neurons (magenta) and epidermal cells in HY and LY conditions in 2D projections. The insets for (A) and (B) show FoxO-GFP levels at ddaC somas (1) and epidermal cells (2) in single confocal sections. (C) Quantification of FoxO-GFP intensity in ddaC neuron soma and epidermal cells in HY and LY conditions. Each circle represents a segment; epi: n=18 for HY, n=14 for LY; ddaC: n=17 for HY and n=14 for LY. (D and E) *Gal4^foxo^*-driven *tdTom* (magenta) in da neurons (green) and epidermal cells in HY and LY conditions in 2D projections. The insets in (D) and (E) show *Gal4^foxo^*-driven *tdTom* expression levels at the soma of ddaC (1) and epidermal cells (2) in single confocal sections. (F) Quantification of *Gal4^foxo^*-driven *tdTom* intensity in ddaC neuron soma and epidermal cells in HY and LY conditions. Each circle represents a segment; epi: n=19 for HY, n=19 for LY; ddaC: n=18 for HY, n=18 for LY. (G-L) ddaC neurons in the *Gal4^ppk^* control (G and J) and animals expressing *Gal4^ppk^*-driven *foxo* RNAi (H and K) and *foxo* (I and L) in HY and LY conditions. (G) and (J) are the same as Figure 2A and 2D, respectively. (M) Quantification of normalized dendritic length in control, *foxo* RNAi and *foxo* overexpression animals in HY and LY conditions. Each circle represents a neuron; HY: n=17 for *Gal4^ppk^*, n=11 for *foxo* RNAi, n=12 for *foxo* OE; LY: n=15 for *Gal4^ppk^*, n=13 for *foxo* RNAi, n=12 for *foxo* OE. *Gal4^ppk^* is the same dataset as figure 2G. (N-Q) Epidermal cells in the *Gal4^R16D01^* control and animals expressing *Gal4^R16D01^*-driven *Atg8a* RNAi in HY and LY conditions. (N) and (P) are the same as Figure 3D and 3F, respectively. (R) Quantification of epidermal cell size ratio (*Gal4^R16D01^* cells / WT cells) in HY and LY conditions. Each circle represents a segment; HY: n=18 for control, n=10 for *foxo* RNAi; LY: n=16 for control, n=18 for *foxo* RNAi. For all quantifications, ***p<0.001; **p<0.01; *p<0.05; ns, not significant; One-way ANOVA and Tukey’s HSD test. Black bars, mean; red bars, SD. Scale bars, 10 μm in (A), (B), (D) and (E); 100 μm in (G-L); 50 μm in (N-Q).

Loss-of-function (LOF) and gain-of-function (GOF) studies of *foxo* previously carried out in C4da neurons showed that FoxO plays a role in enhancing dendritic space-filling by stabilizing dendritic microtubule (Sears and Broihier, 2016). However, how nutrients may influence FoxO’s function is not clear. We therefore knocked down and overexpressed FoxO in C4da neurons in the HY and LY media. Expressing *foxo-RNAi* in epidermal cells effectively eliminated a co-expressed FoxO-GFP (Figures S4G-S4I), confirming the effectiveness of this RNAi line. Knocking down *foxo* in C4da neurons did not affect normalized dendrite length in either HY or LY condition (Figures 4G, 4H, 4J, 4K and 4M), but caused 13% reduction of dendrite density only in LY medium. For unknown reasons, the dendritic fields of *ppk*>*foxo-RNAi* neurons were larger than those of WT controls under LY, contributing to the decrease of dendrite density of *foxo* knockdown neurons despite at similar dendrite lengths. On the other hand, overexpressing FoxO in C4da neurons caused mild (11.7%) and strong (49.2%) reduction of normalized dendrite length in HY and LY conditions, respectively (Figures 4I, 4L, and 4M). We next examined the role of FoxO in epidermal cell growth by knocking down *foxo* using *Gal4^R16D01^*. The knockdown had no effect in HY, but increased the epidermal cell size by 34% under the LY condition (Figures 4N-4R). Above results collectively suggest that FoxO affects the growth of C4da neurons and epidermal cells differentially: In neurons, although high FoxO levels are growth-inhibitory, endogenous FoxO is likely expressed at too low levels to directly affect dendrite growth; in epidermal cells, FoxO is expressed at a much higher level, but only inhibit cell growth under nutrient stress.

**Figure S4.**
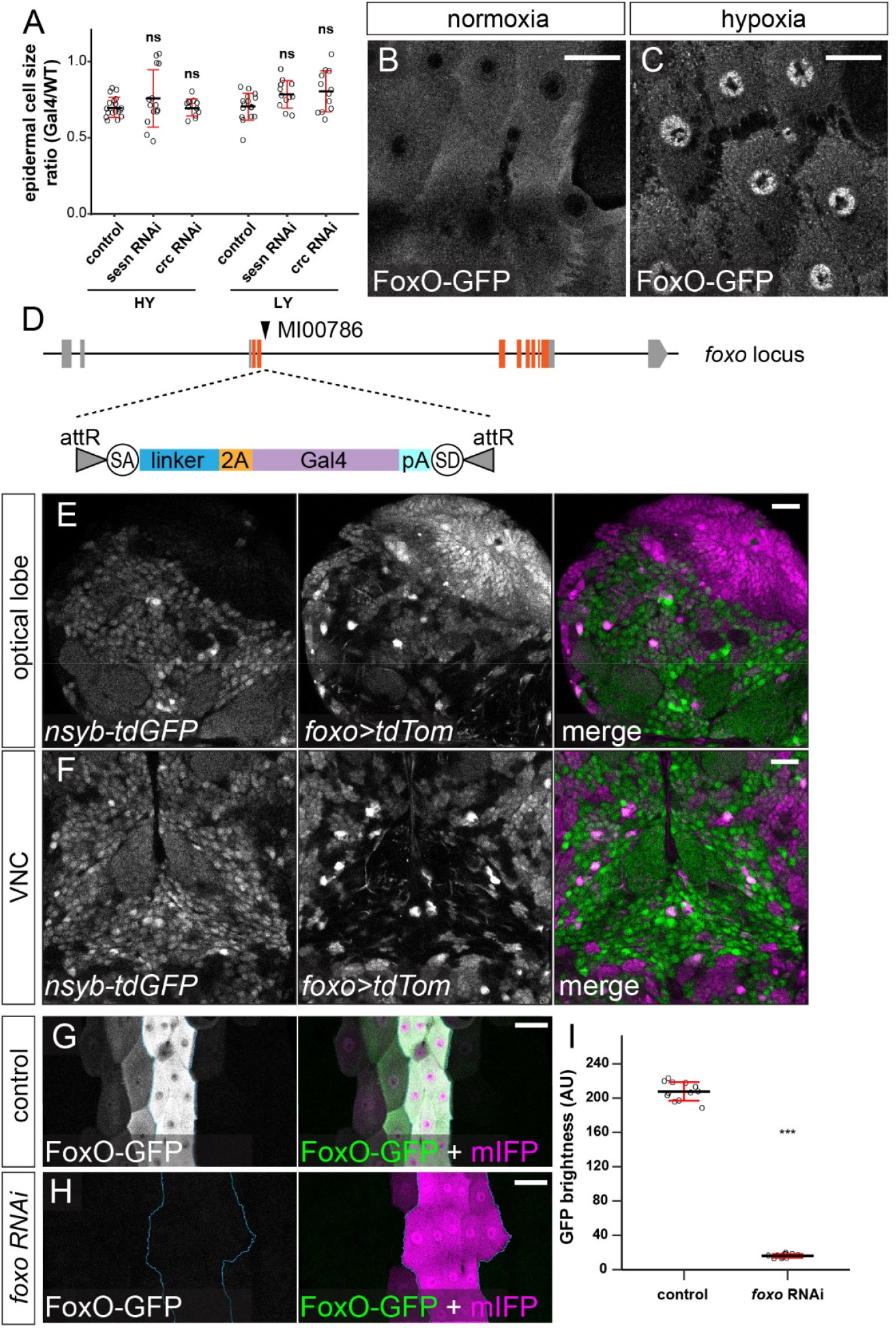
Location of FoxO and effectiveness of *foxo* RNAi. (A) Quantification of epidermal cell size ratio (*Gal4^R16D01^* cells / WT cells) in HY and LY conditions for genotypes indicated. Each circle represents a segment; HY: n=18 for control, n=14 for *sesn* RNAi, n=12 for *crc* RNAi; LY: n=16 for control, n=12 for *sesn* RNAi, n=12 for *crc* RNAi. (B and C) FoxO-GFP expression in epidermal cells under normoxia (B) and hypoxia (C). (D) Diagram of FoxO-Gal4. (E and F) *Gal4^foxo^*-driven *tdTom* expression in optical lobe (E) and VNC (F) in LY condition. (G and H) FoxO-GFP expression in control (G) and *Gal4^R16D01^*-driven *foxo* RNAi animals. Blue line in (H) outlines the *Gal4^R16D01^* domain. (I) Quantification of FoxO-GFP levels in control and *foxo* RNAi animals. Each circle represents a segment; n=12 for control and *foxo* RNAi. ***p<0.001; Student’s t-test. Black bars, mean; red bars, SD. Scale bars, 25 μm in (B) and (C), 20 μm in (E) and (F), 50 μm in (G) and (H).

### Overexpressed FoxO exerts its effects through modulating Tor signaling and autophagy

We next used pRpS6 staining to examine whether overexpressed FoxO affects dendrite growth by inhibiting Tor signaling. In HY, neuronal FoxO overexpression did not cause detectable changes in the ratio of pRpS6 levels between neuronal compartments and epidermal cells (Figures 2M, 5A, 5C, and 5D). However, in LY, while pRpS6 levels in the somas of FoxO-overexpressing neurons were still higher than in epidermal cells, the dendritic pRpS6 signal was reduced to levels lower than those of epidermal cells (Figures 5B-5D). We further examined the autophagy level of FoxO-overexpressing neurons using mCherry-Atg8a. While the autophagosome level in C4da cell bodies was not altered by FoxO-expression in HY, it increased 4.9 folds in LY (Figures 4E-4G). These data suggest that nutrient stress enables overexpressed FoxO to suppress dendritic Tor signaling and induces autophagy in neurons. The lack of high FoxO expression in wildtype neurons thus ensures preferential dendrite growth in nutrient stress by protecting dendritic Tor signaling and suppressing autophagy.

**Figure 5.**
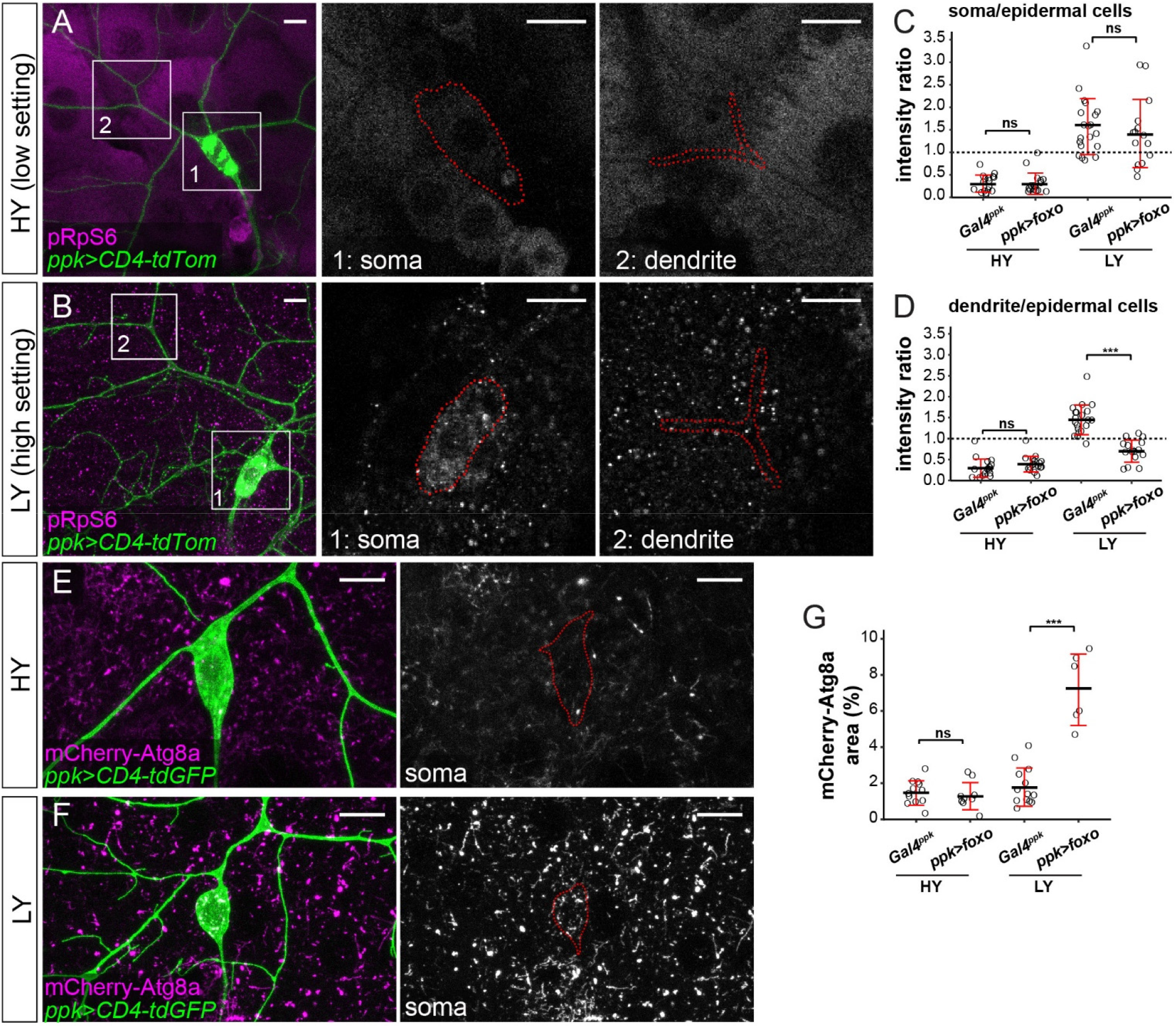
Overexpressed FoxO exerts its effects through modulating Tor signaling and autophagy. (A and B) pRpS6 staining (magenta) of ddaC neurons (green) and epidermal cells in *Gal4^ppk^*-driven *UAS-foxo* animals in HY and LY conditions in 2D projections. The insets for (A) and (B) show pRpS6 staining at the soma (1) and primary dendrites (2) in single confocal sections, with the somas and dendrites outlined. (C and D) Quantification of pRpS6 intensity ratios in control and *foxo* OE animals in HY and LY conditions. Each circle represents a segment; HY: n=17 for control, n=17 for *foxo* RNAi; LY: n=20 for control, n=16 for *foxo* RNAi. The control datasets are the same as in Figure 2P. (E and F) mCherry-Atg8a (magenta) in ddaC soma (green) of *Gal4^ppk^*-driven *UAS-foxo* animals in HY and LY. The soma images are projections from thinner volumes only containing the soma. (G) Quantification of mCherry-Atg8a levels in ddaC soma measured by the area percentage of mCherry-Atg8a-positive vesicles. Each circle represents a neuron; HY: n=12 for control, n=9 for *foxo* OE; LY: n=13 for control, n=6 for *foxo* OE. For all quantifications, ***p< 0.001; ns, not significant; One-way ANOVA and Tukey’s HSD test. Black bars, mean; red bars, SD. Scale bars, 10 μm.

### Nutrient stress-induced dendrite overgrowth sensitizes neurons

C4da neurons are polymodal nociceptive neurons that respond to extreme stimuli such as heat, mechanical forces, and UV light (Tracey et al., 2003; Hwang et al., 2007; Xiang et al., 2010). To test whether the preferential dendrite growth under nutrient stress is physiologically relevant, we examined the function of C4da neurons using an established heat-response assay (Babcock et al., 2009). In this assay, a temperature-controlled heat probe is used to elicit a C4da-dependent larval rolling behavior within 5 s (fast response) or 20 s (response) of heat stimulus. We first examined wildtype larvae reared in HY and LY media. Interestingly, at a given temperature, a larger percentage of LY larvae showed response (from 40°C to 46°C) and fast response (from 45°C to 48°C) than HY larvae (Figures 6A and 6B). In general, the temperatures required to induce response in a similar percentage of larvae were generally 2°C lower for the LY condition than for the HY condition. When tested specifically at 46°C, most of the LY larvae responded within 6 s while HY larvae showed a broader range of durations needed to respond (Figure 6C). These data suggest that nutrient stress sensitizes larvae so that they react more acutely to noxious heat.

**Figure 6.**
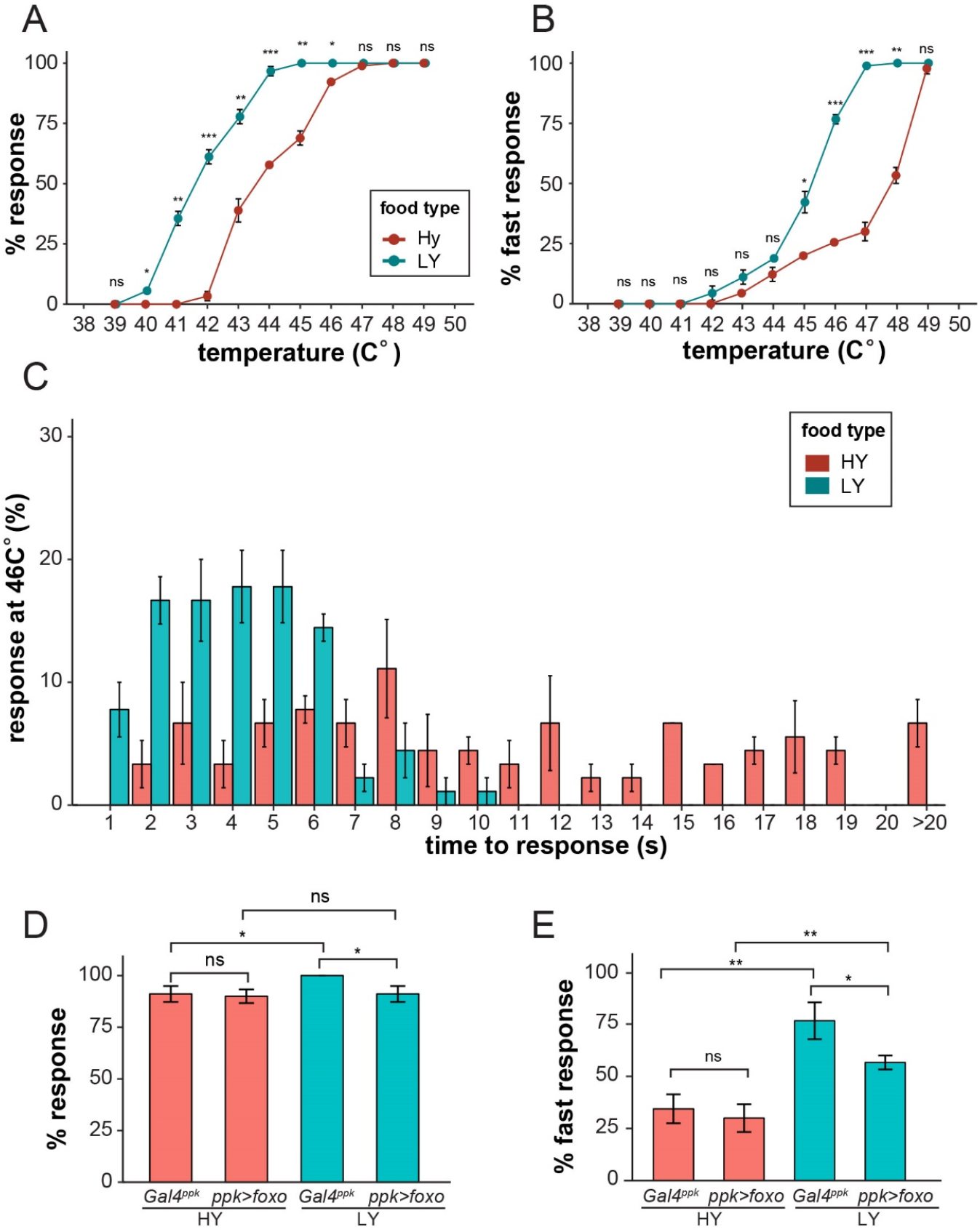
Dendrite overgrowth under nutrient stress sensitized neurons. (A) A plot showing the percent of responders (respond within 20s) versus temperature. n = number of larvae; n=90 for HY and LY at each temperature. (B) A plot showing the percent of fast responders (respond within 5s) versus temperature. n = number of larvae; n=90 for HY and LY at each temperature. (C) A plot showing the percent of responders at 46°C versus the response time. n = number of larvae; n=90 for HY and LY. In all panels, error bars indicate the standard error (SE). (D) A plot showing the percent of responders versus temperature for *Gal4^ppk^* and FoxO OE animals in HY and LY conditions at 46°C. n=number of larvae; n=120 for each group. (E) A plot showing the percent of fast responders versus temperature for *Gal4^ppk^* and FoxO OE animals in HY and LY conditions at 46°C. n=number of larvae; n=120 for each group. For all quantifications, ***p< 0.001; **p<0.01; *p<0.05: ns, not significant; Student’s *t* test.

We next examined the effects of FoxO overexpression in C4da neurons by stimulating the larvae at 46°C. FoxO overexpression did not significantly change the rolling behviors of HY larvae but reduced percentages of LY larvae that showed response and fast response (Figures 6D and 6E). This decrease of nociceptive response thus correlates with the strong dendrite reduction of FoxO-overexpressing neurons under nutrient restriction.

### Different types of somatosensory neurons respond to nutrient stress differentially

Among the four classes of da neurons, C4da neurons exhibit space-filling and are highly dynamic (Grueber et al., 2003; Poe et al., 2017), while C1da and C2da neurons have simple arbors and sparse dendrites occupying defined territories and grow mostly by scaled expansion of the existing dendritic arbors established in late embryogenesis (Grueber et al., 2002). C3da neurons grow more complex dendritic arbors and are characterized by numerous short terminal branches called dendritic spikes, which are highly dynamics during larval growth (Grueber et al., 2002; Nagel et al., 2012). We asked whether nutrient stress also impacts dendritic growth of C1da and C3da neurons. Consistent with the previous report by Watanabe et al. (Watanabe et al., 2017), the yeast concentration did not have obvious effects on the total dendrite length of C1da neuron ddaE (Figures 7A-7C). However, nutrient stress stimulated the dendritic growth of C3da neurons ddaA and ddaF: the total dendrite length increased by 40% and 32%, respectively; the total terminal dendrite length increased by 61% and 46%, respectively; the terminal branch numbers increased by 47% and 58%, respectively (Figures 7D-7H). These data suggest that nutrient stress promotes overbranching of complex dendritic arbors of C3da and C4da neurons but not simple arbors of C1da neurons.

**Figure 7.**
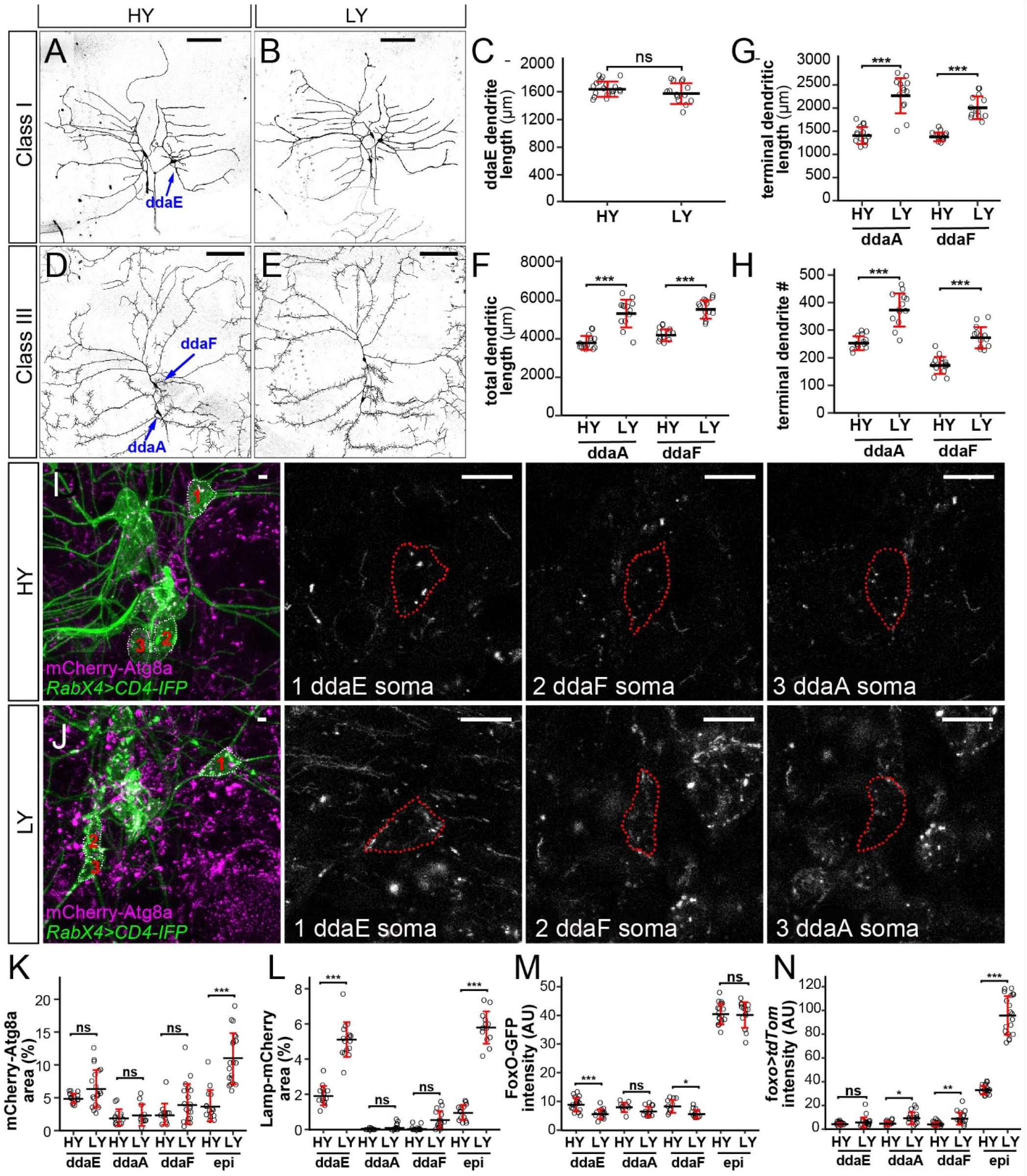
Different types of somatosensory neurons respond to nutrient stress differentially. (A and B) ddaE neurons in HY and LY conditions. (C) Quantification of ddaE dendrite length in HY and LY conditions. (D and E) ddaF and ddaA neurons in HY and LY conditions. (F-H) Quantification of total dendritic length (F), terminal dendritic length (G) and terminal dendrite number (H) of ddaF and ddaA neurons in HY and LY conditions. (I and J) Autophagy levels (magenta) in da neuron soma (green) and epidermal cells in HY and LY conditions. Autophagy levels are measured by the area of mCherry-Atg8a-positive vesicles. 1, 2 and 3 show autophagy levels of ddaE, ddaF and ddaA soma outlined by white dots in the merge channel. Red dots outline the soma of da neurons. (K and L) Quantification of autophagy levels in da neuron soma and epidermal cells in HY and LY conditions measured by the the area of mCherry-Atg8a-positive vesicles (K) and Lamp-mCherry-positive vesicles (L). (M and N) Quantification of FoxO expression level (M) and *Gal4^foxo^*-driven *tdTom* expression level (N) in da neuron soma in HY and LY conditions. For all quantifications, ***P< 0.001; **P<0.01; *P<0.05: ns, not significant; ANOVA and Tukey’s HSD test. Each circle represents a neuron. Black bars, mean; red bars, SD. Scale bars, A, B, D and E 100um; I and J 10um.

We then examined whether C1da and C4da neurons are subjected to the same regulation of autophagy as C4da neurons under nutrient stress. Interestingly, the cell bodies of these neurons showed variable and cell-specific baseline autophagy levels in HY, as indicated by mCherry-Atg8a, but these levels did not appear to be altered by the nutrient level (Figures 7I-7K). Similarly, C3da neurons showed low and nutrient-independent lysosomal level (Figure S5A, S5B, 7C). Interestingly, the C1 ddaE showed a much higher level of the lysosomal marker Lamp-mCherry than other da classes in HY, and this level was enhanced 2.7 folds by nutrient stress (Figure S5A, S5B, 7C), suggesting that C1da neurons have uniquely high and nutrient-dependent lysosomal system. Lastly, we examined *foxo* expression in C1da and C3da neurons using *foxo-GFP* and *foxo-Gal4*. Similar to C4da neurons, C1da and C3da neurons showed only background-noise levels of FoxO-GFP and*foxo*>*tdTom* signals (Figures 7M and 7N). These data suggest that, similar to C4da neurons, C1da and C3da neurons also exhibit low FoxO expression and nutrient-independent autophagy levels, even though only C3da neurons, but not C1da neurons, show nutrient stress-induced dendrite overgrowth.

**Figure S5.**
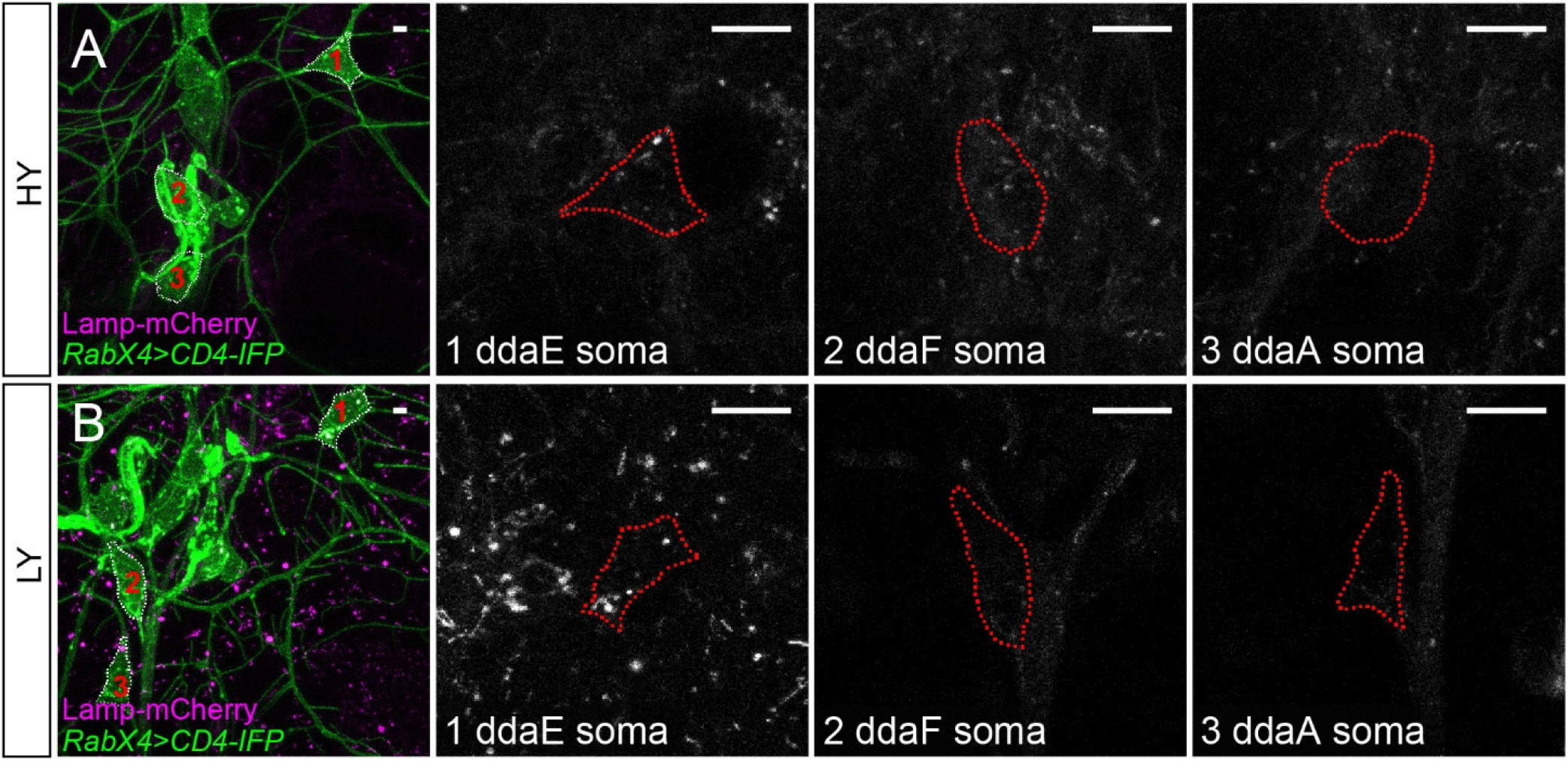
Autophagy levels in ddaC neurons and epidermal cells. (A and B) Autophagy levels (magenta) in da neuron soma (green). Autophagy levels are measured by the area of Lamp-mCherry-positive vesicles. 1, 2 and 3 show autophagy levels of ddaE, ddaF and ddaA soma outlined by white dots in the merge channel. Red dots outline the soma of da neurons. Scale bars, 10um.

## DISCUSSION

In this study, we show that *Drosophila* C3da and C4da neurons exhibit a growth advantage over neighboring epidermal cells under nutrient restriction, resulting in dendrite overgrowth. This growth advantage is made possible by the differential FoxO expression and autophagic responses of neurons and non-neural cells (Figure 8). In non-neural tissues like epidermal cells, the stress sensor FoxO and the master regulator of autophagic genes Mitf are amply expressed, allowing epidermal cells to respond robustly to nutrient stress. In these tissues, high nutrition elevates InR/Tor signaling and suppresses FoxO activity and autophagy, leading to a high growth rate; in low nutrients, the reduction in InR/Tor signaling combined with high FoxO activity stimulates autophagy and greatly slows down the increase in cell size. In contrast, PNS neurons express very little FoxO and, likely due to the absence of Mitf expression, exhibit low basal levels of autophagy. As a result, the dendritic InR/Tor signaling is not further suppressed by FoxO when the systemic insulin level is low. Therefore, the low FoxO expression and the lack of autophagy induction in PNS neurons protect dendrite growth from nutrient stress.

**Figure 8.**
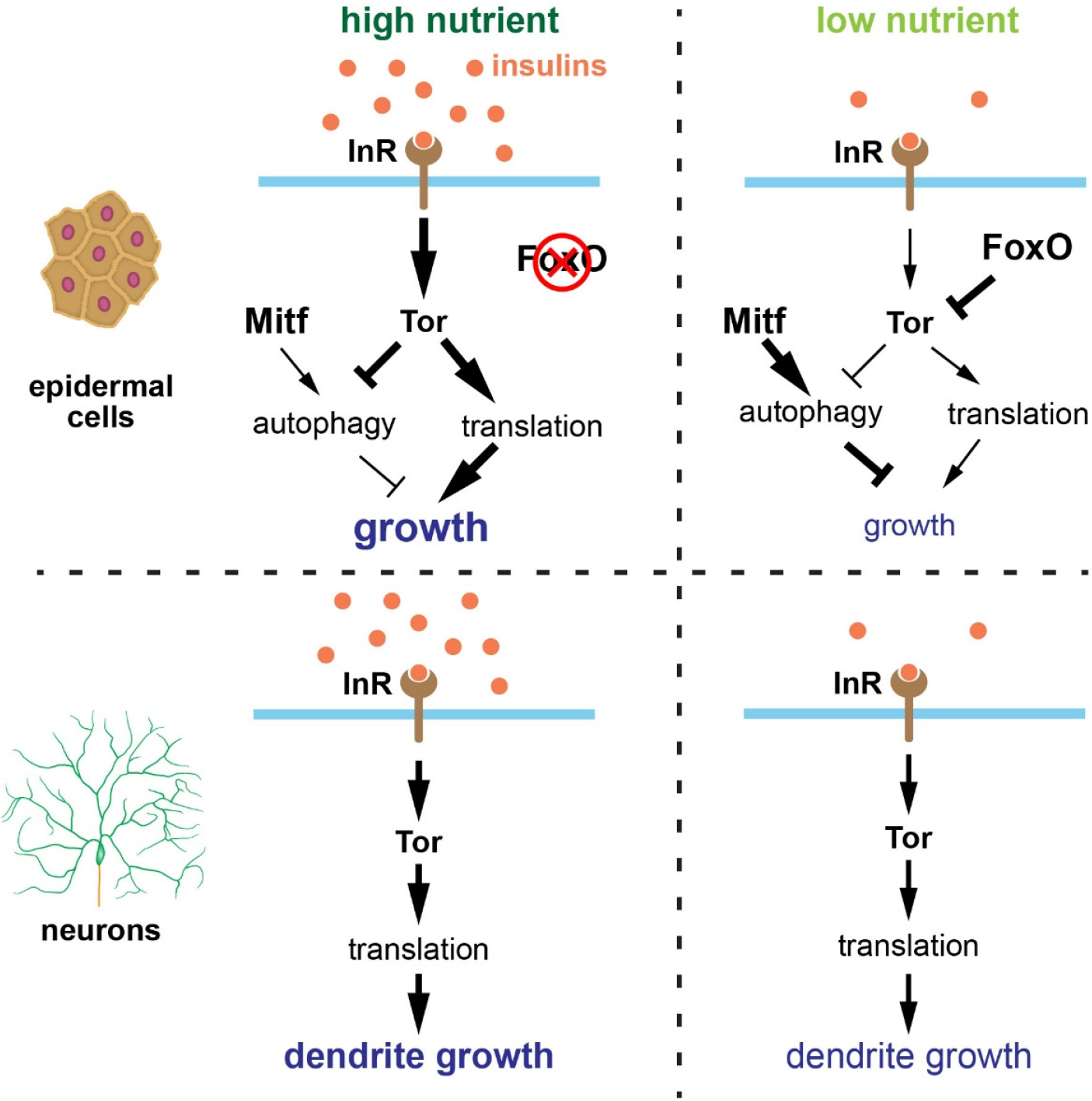
Model of da neurons and epidermal cells growth control.

Brain sparing has been recognized in both mammals and insects as an important means to protect the developing nervous system from nutrient deficiency. The preferential dendrite growth of da neurons constitutes another form of nervous system sparing that bears important distinctions from *Drosophila* brain sparing. First, while the CNS is protected against starvation in the proliferation of neural stem cells (Cheng et al., 2011), post-mitotic da neurons are spared at the level of neuronal arbor growth of individual cells. Second, unlike CNS neuroblasts, which rely on a special extrinsic factor (the glial-derived Jeb) to sustain neurogenesis, PNS neurons possess a unique intrinsic genetic program (in FoxO and autophagic gene expression) that endows them the resistance to nutrient stress. Lastly, the CNS sparing is independent of InR and Tor, made possible by the alternative Jeb/Alk/PI3K pathway, while the PNS protection still relies on InR/Tor signaling. Therefore, our work reveals a novel mechanism of neural protection under nutrient stress.

FoxO proteins have been demonstrated to play important roles in the nervous system. In the mammalian brain, FoxO members are highly expressed in NSCs and are required for the long-term maintenance of the adult NSC pool critical for adult neurogenesis (Paik et al., 2009; Renault et al., 2009; Yeo et al., 2013). Recently, FoxOs were also found to regulate dendrite branching and spine density of adult-born hippocampal neurons (Schaffner et al., 2018). Interestingly, FoxOs in these neurons suppress mTor signaling and maintain a level of autophagic flux that is necessary for the normal morphogenesis of the neurons. In addition, FoxOs were found to be upregulated in aged brains and function to delay aging-related axonal tract degeneration by suppressing mTor activity (Hwang et al., 2018). In *Drosophila* C4da neurons, FoxO was previously found to promote dendrite space-filling and to mediate polyQ-induced neuronal toxicity (Sears and Broihier, 2016; Kwon et al., 2018). However, FoxO expression patterns in neural and non-neural tissues have not been compared and whether neuronal function of FoxO is related to nutrition was previously unknown. Our results show that FoxO is expressed at a much lower level in da neurons than in non-neural larval tissues. Consequently, inhibiting *foxo* in neurons does not directly affect dendrite growth, even though high Foxo levels in neurons inhibits dendrite growth. Nevertheless, our results support that neuronal FoxO mildly promotes dendritic space-filling of C4da neurons only under nutrient restriction by reducing the size of the dendrite field though an unknown mechanism. More importantly, the lack of high FoxO expression makes neurons insensitive to nutrient stress, giving them a growth advantage over non-neural tissues that express higher levels of FoxO. Interestingly, this FoxO-dosage-dependent nutrient-insensitivity has been previously described in the adult genitalia (Tang et al., 2011). While the adult wings and maxillary palps become small on poor diets, the genital arches show much weaker response. The differential responses are linked to tissue-intrinsic levels of FoxO expression. However, in this example and previously described FoxO functions in growth regulation, FoxO mostly regulates cell numbers but not cell size (Junger et al., 2003; Puig et al., 2003). Therefore, our study reals a novel function for FoxO in environmental regulation of neural development. It’s worth noting that although FoxO is minimally expressed in da neurons, it is expressed at a higher level in a subpopulation of CNS neurons, raising the possibility that the arbor growth of these neurons may be differentially regulated by nutrient availability.

Lastly, our results reveal a level of neuronal diversity in the response to nutrient stress. Although all da neurons we examined show similarly low FoxO expression and the lack of autophagy induction under nutrient restriction, only class III and IV but not class I neurons display preferential dendrite growth. This distinction may be related to their arbor growth mechanisms and neuronal functions. As proprioceptive neurons that detect body surface folds during locomotion (He et al., 2019; Vaadia et al., 2019), C1da neurons have simple arbors covering defined territories on the larval body wall. Their dendritic arbors grow mainly by expanding the shafts of the dendritic branches established during embryogenesis. Their functional demands may require a tighter growth coupling with the epidermis but not with environmental nutrition availability. In contrast, C3da and C4da neurons have highly dynamic high-order branches that grow also by branching and tip extension. The lack of FoxO-mediated growth suppression therefore leads to dendrite hyperarborization. As these neurons sense mild to extreme levels of external stimuli, the heightened sensations allowed by their dendrite overgrowth may bestow the larva a survival advantage in an unfavorable environment.

## METHODS AND MATERIALS

### Key Resources Table

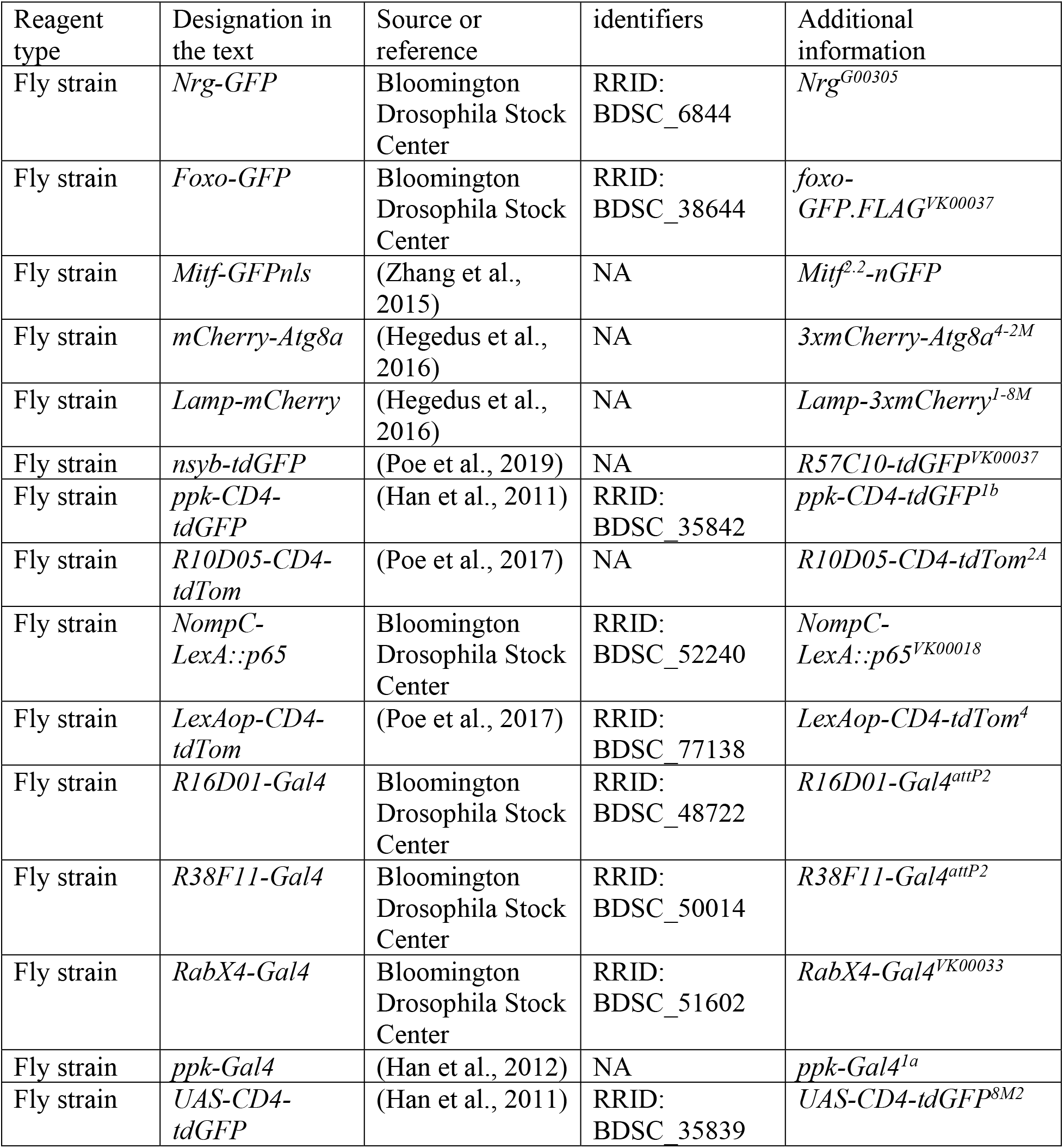
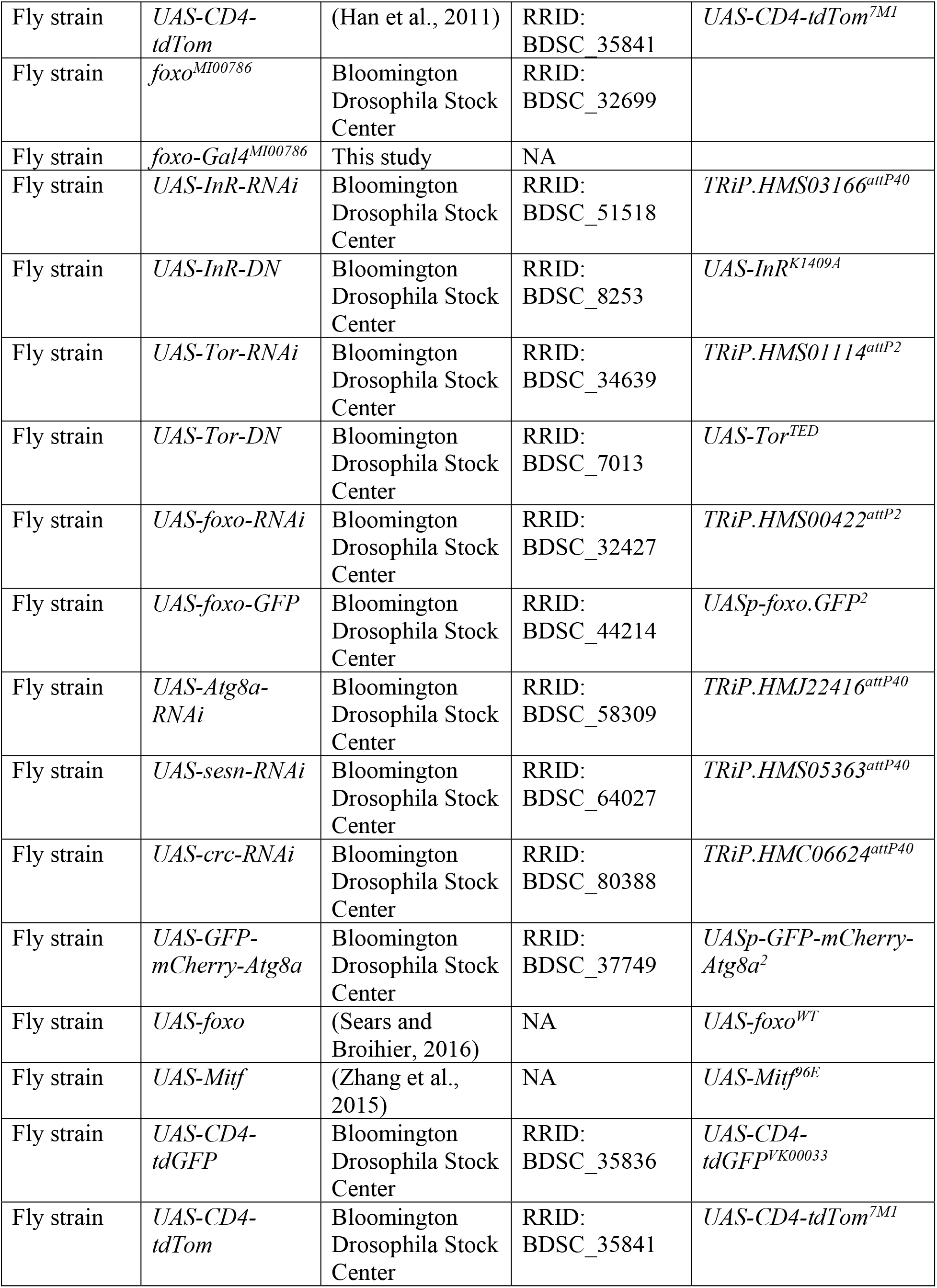
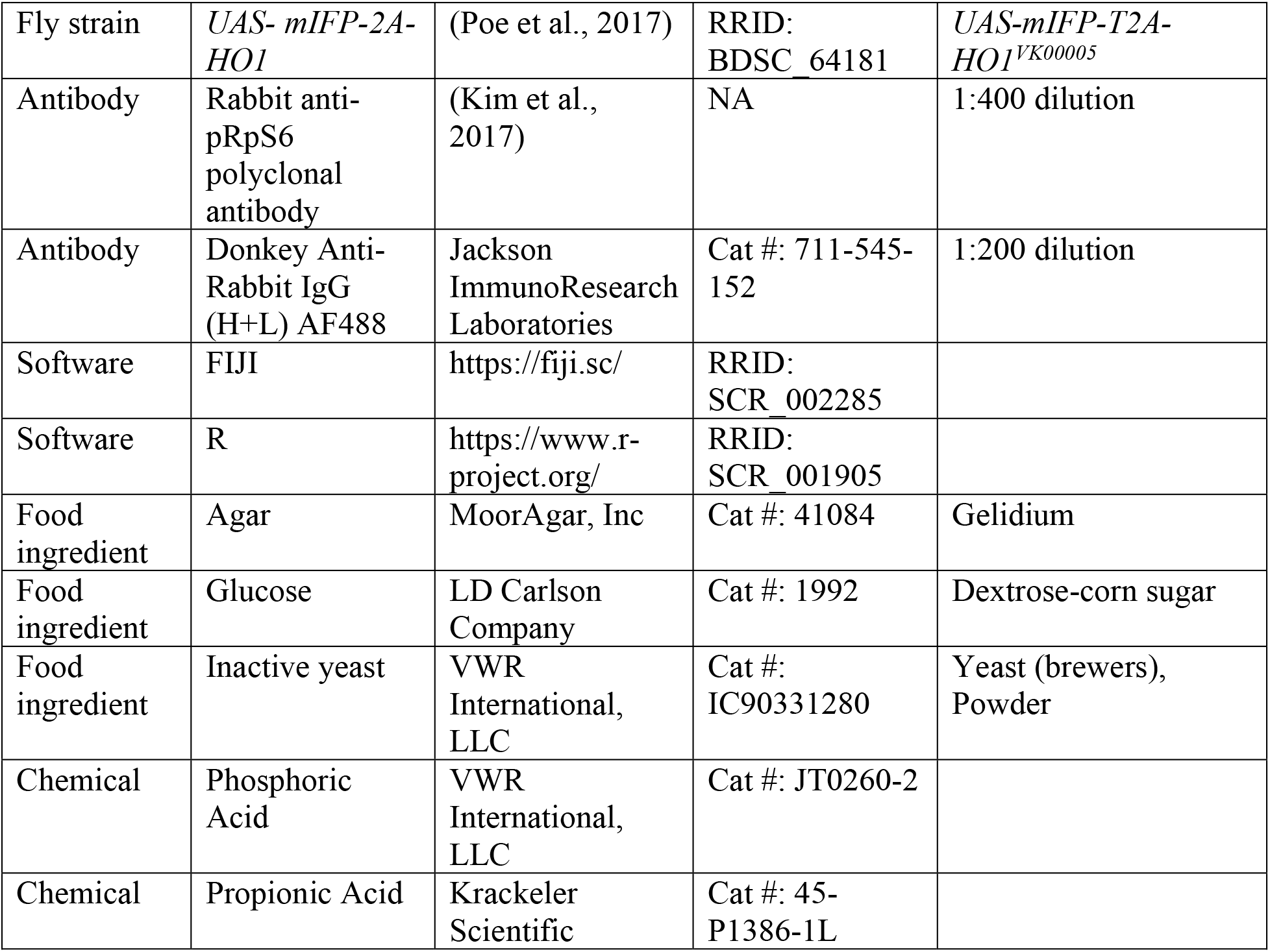

### Live imaging

Animals were raised at 25°C in density-controlled vials containing between 50-70 embryos collected in a 3-hour time window. To achieve optimum embryo densities, approximately 50 virgin females were aged 5 days on molasses food with yeast paste, crossed with approximately 15-20 males, and then allowed to mate for 1-2 days on molasses food with yeast paste. Embryo collections were then performed in a 3-hour time window on both 1% and 8% yeast food. Third instar larvae at 86 hrs AEL on 8% yeast or 216 hrs AEL on 1% yeast were mounted in glycerol and imaged with a Leica SP8 confocal. The A2-A3 segments of 8-10 larvae were imaged for each genotype using a 20X oil objective. To image larvae younger than 72 hrs AEL, larvae were anesthetized by isoflurane for 2 minutes & then mounted in halocarbon oil. To image FoxO-GFP under normoxia, larvae were handled with care and imaged within 2 minutes after mounting on slides. For imaging under hypoxia, larvae were left on the slides for 5 minutes before imaging.

### Fly Food Recipe

Fly food was prepared using the following recipes (for the dispersal of ~12 mL into 20 vials).

|  |  |  |
| --- | --- | --- |
| Ingredients: | 1% yeast | 8% yeast |
| Distilled H<sub>2</sub>O | 240mL | 234mL |
| Agar | 2.4g (12g/L) | 2g (10g/L) |
| Glucose | 20g | 20g |
| Inactive yeast | 2.5g | 20g |
| Acid mix (phosphoric acid + propionic acid) | 2mL | 2mL |
| Target final solution volume | 250mL | 250mL |

Acid Mix was made by preparing Solution A (41.5 ml Phosphoric Acid mixed with 458.5 ml distilled water) and Solution B (418 ml Propionic Acid mixed with 82 ml distilled water) separately and mixing Solution A and Solution B together.

### Fly Stocks

The strains used in this study are listed in the Key Resources Table. We used the following neuronal markers to label specific classes of da neurons: *ppk-CD4-tdGFP*, *ppk-Gal4*, *UAS-CD4-tdGFP* and *UAS-CD4-tdTom* for C4da; *R10D05-CD4-tdTom* for C1da; *NompC-LexA::p65 LexAop-CD4-tdTom* for C3da; *RabX4-Gal4 UAS-mIFP-2A-HO1* for all classes of da neurons. *ppk-Gal4* was used for RNAi knockdown and overexpression in C4da neurons. *Gal4^R38F11^* and *Gal4^R16D01^* are used for RNAi knockdown and overexpression in all or stripes of epidermal cells, respectively.

*foxo-Gal4[MI00786]* was generated using Trojan-MiMIC system (Diao et al., 2015). MiMIC line MI00786 was crossed to flies with the triplet Trojan donor construct. The progeny of this cross were then crossed to females expressing Cre recombinase and ΦC31 integrase in the germline which allow the Trojan exons to replace the MiMIC attP cassettes. Progeny were then crossed to *UAS-GFP* for screening Gal4 expression by fluorescence microscopy. Adults positive for GFP expression were used to establish the line. The 2A-Gal4 insertions from established lines were sequenced to confirm the accuracy of the site and the reading frame.

### Image quantification

#### Neuron quantification

Unless noted otherwise, only larvae with the segment width falling between 500-550 μm were quantified for neurons. The tracing and measurement of da neuron dendrites were done in Fiji/ImageJ as previously described (Poe et al., 2017). Briefly, images of dendrites (1,024 × 1,024 pixels) taken with a 20X objective were first processed sequentially by Gaussian Blur, Auto Local Threshold, Particles4, Skeletonize (2D/3D), and Analyze Skeleton (2D/3D) plugins. The length of skeletons was calculated based on pixel distance. Dendrite density was calculated using the formula: 1000 × dendritic length (μm)/dendritic area (μm^2^); normalized dendrite length was calculated as dendritic length (μm)/segment width (μm). Normalized length ratio was calculated using the formula: normalized dendrite length on LY/normalized dendrite length on HY.

#### Epidermal cell quantification

Images of epidermal cells labeled by Nrg-GFP (1,024 × 1,024 pixels) taken with a 20X objective were first processed by Gaussian Blur (Sigma: 1) and then Auto Local Threshold (Phansalkar method, radius: 30). Isolated particles below the size of 500 pixels were removed by the Particles4 plugin (http://www.mecourse.com/landinig/software/software.html). The Nrg-GFP signal was then converted to single-pixel-width skeletons of epidermal cell borders using the Skeletonize (2D/3D) plugin. Images were then visually inspected to ensure that all epidermal cell borders were accurately labeled. Any erroneous epidermal cell borders were removed. Regions of interest (ROIs) were manually drawn to encompass the epidermal cells for quantification. Analyze Particles was then used to measure area, perimeter, height (Feret), and width (minFeret) for each epidermal cell in the ROIs. Epidermal cell size ratio was calculated as average cell size in the RNAi-expressing region/average cell size in the WT region.

#### Other image quantification

For quantification of Lamp-mCherry and Atg8a-mCherry in Fiji/ImageJ, Z-stack images of dendrites and Lamp-mCherry or Atg8a-mCherry (1,024 × 1,024 pixels) taken with a 40X objective and a step size of 0.5 μm were converted into binary masks using thresholding. For quantification in epidermal cells, the stacks were projected into 2D images. ROIs were drawn manually outside the neuron. For quantification in neurons, care was taken to select the optical sections only containing the cell body but not Lamp-mCherry or Atg8a-mCherry signals below or above. The optical sections were then projected into 2D images, and ROIs covering cell bodies were generated based on masks of the dendrite channel. Finally, the mask area percentage within each ROI was calculated.

Images of *Mitf-GFPnls* (1,024 × 1,024 pixels) were taken with a 40X objective. A ROI on neuronal or epidermal cell nucleus was drawn manually on a single slice with the strongest signal. The mean gray value of the area was calculated.

Autophagic flux levels were measured by Atg8a-mCherry area percentage divided by GFP intensity. Images of epidermal cells and ddaC neurons (1,024 × 1,024 pixels) were taken with a 40X objective and a step size 0.5 μm. Atg8a-mCherry area percentages were determined as described above. For GFP intensity, a ROI in ddaC neuron cell body or epidermal cells were drawn manually using the maximum projected image and the mean gray value of the area was calculated.

### Immunostaining

Antibody staining was done as previously described (Poe et al., 2017). Briefly, third instar larvae were dissected in cold PBS, fixed in 4% formaldehyde/PBS for 20 min at room temperature, and stained with the proper primary antibodies and subsequent secondary antibodies, each for 2 hrs at room temperature. The details of the antibodies used are in the Key Resources Table.

### Behavior assay

Larval heat-induced pain response was measured as described previously (Babcock et al., 2009). Wandering third larvae were scooped out of the food and gently cleaned with water, then transferred into a small petri dish with water drops to keep the animals moist. A temperature-controlled heat probe (ProDev Engineering, TX) was used to apply the heat onto the larval body surface. The stimulus was delivered by gently touching the animals laterally on segment A4. Each animal can only be tested once. The response latency was measured from the start of touch on the animal until it initiated a complete 360° roll.

## ACKNOWLEDGMENTS

We thank Francesca Pignoni, Gábor Juhász, and Bloomington Stock Center for fly stocks; Jongkyeong Chung for antibody; Michael Goldberg for critical reading and suggestions on the manuscript. This work was supported by a Cornell start-up fund and NIH grants (R01NS099125 and R21OD023824) awarded to C.H.

## REFERENCES

Babcock, D.T., Landry, C., and Galko, M.J., 2009. Cytokine signaling mediates UV-induced nociceptive sensitization in Drosophila larvae. Curr Biol 19, 799–806. https://doi.org/10.1016/j.cub.2009.03.062.

Boulan, L., Milan, M., and Leopold, P., 2015. The Systemic Control of Growth. Cold Spring Harb Perspect Biol 7. https://doi.org/10.1101/cshperspect.a019117.

Brogiolo, W., Stocker, H., Ikeya, T., Rintelen, F., Fernandez, R., and Hafen, E., 2001. An evolutionarily conserved function of the Drosophila insulin receptor and insulin-like peptides in growth control. Curr Biol 11, 213–221.

Burnett, P.E., Barrow, R.K., Cohen, N.A., Snyder, S.H., and Sabatini, D.M., 1998. RAFT1 phosphorylation of the translational regulators p70 S6 kinase and 4E-BP1. Proc Natl Acad Sci U S A 95, 1432–1437. https://doi.org/10.1073/pnas.95.4.1432.

Cheng, L.Y., Bailey, A.P., Leevers, S.J., Ragan, T.J., Driscoll, P.C., and Gould, A.P., 2011. Anaplastic lymphoma kinase spares organ growth during nutrient restriction in Drosophila. Cell 146, 435–447. https://doi.org/10.1016/j.cell.2011.06.040.

Diao, F., Ironfield, H., Luan, H., Diao, F., Shropshire, W.C., Ewer, J., Marr, E., Potter, C.J., Landgraf, M., and White, B.H., 2015. Plug-and-play genetic access to drosophila cell types using exchangeable exon cassettes. Cell Rep 10, 1410–1421. https://doi.org/10.1016/j.celrep.2015.01.059.

Eijkelenboom, A., and Burgering, B.M., 2013. FOXOs: signalling integrators for homeostasis maintenance. Nat Rev Mol Cell Biol 14, 83–97. https://doi.org/10.1038/nrm3507.

Fullgrabe, J., Ghislat, G., Cho, D.H., and Rubinsztein, D.C., 2016. Transcriptional regulation of mammalian autophagy at a glance. J Cell Sci 129, 3059–3066. https://doi.org/10.1242/jcs.188920.

Ganley, I.G., Lam du, H., Wang, J., Ding, X., Chen, S., and Jiang, X., 2009. ULK1.ATG13.FIP200 complex mediates mTOR signaling and is essential for autophagy. J Biol Chem 284, 12297–12305. https://doi.org/10.1074/jbc.M900573200.

Geminard, C., Rulifson, E.J., and Leopold, P., 2009. Remote control of insulin secretion by fat cells in Drosophila. Cell Metab 10, 199–207. https://doi.org/10.1016/j.cmet.2009.08.002.

Grueber, W.B., Jan, L.Y., and Jan, Y.N., 2002. Tiling of the Drosophila epidermis by multidendritic sensory neurons. Development 129, 2867–2878.

Grueber, W.B., Ye, B., Moore, A.W., Jan, L.Y., and Jan, Y.N., 2003. Dendrites of distinct classes of Drosophila sensory neurons show different capacities for homotypic repulsion. Curr Biol 13, 618–626.

Gruenwald, P., 1963. Chronic Fetal Distress and Placental Insufficiency. Biol Neonat 5, 215–265.

Han, C., Jan, L.Y., and Jan, Y.N., 2011. Enhancer-driven membrane markers for analysis of nonautonomous mechanisms reveal neuron-glia interactions in Drosophila. Proc Natl Acad Sci U S A 108, 9673–9678. https://doi.org/10.1073/pnas.1106386108.

Han, C., Wang, D., Soba, P., Zhu, S., Lin, X., Jan, L.Y., and Jan, Y.N., 2012. Integrins regulate repulsion-mediated dendritic patterning of drosophila sensory neurons by restricting dendrites in a 2D space. Neuron 73, 64–78. https://doi.org/10.1016/j.neuron.2011.10.036.

Hattori, Y., Usui, T., Satoh, D., Moriyama, S., Shimono, K., Itoh, T., Shirahige, K., and Uemura, T., 2013. Sensory-neuron subtype-specific transcriptional programs controlling dendrite morphogenesis: genome-wide analysis of Abrupt and Knot/Collier. Dev Cell 27, 530–544. https://doi.org/10.1016/j.devcel.2013.10.024.

He, C., and Klionsky, D.J., 2009. Regulation mechanisms and signaling pathways of autophagy. Annu Rev Genet 43, 67–93. https://doi.org/10.1146/annurev-genet-102808-114910.

He, L., Gulyanon, S., Mihovilovic Skanata, M., Karagyozov, D., Heckscher, E.S., Krieg, M., Tsechpenakis, G., Gershow, M., and Tracey, W.D., Jr., 2019. Direction Selectivity in Drosophila Proprioceptors Requires the Mechanosensory Channel Tmc. Curr Biol 29, 945–956 e943. https://doi.org/10.1016/j.cub.2019.02.025.

Hegedus, K., Takats, S., Boda, A., Jipa, A., Nagy, P., Varga, K., Kovacs, A.L., and Juhasz, G., 2016. The Ccz1-Mon1-Rab7 module and Rab5 control distinct steps of autophagy. Mol Biol Cell 27, 3132–3142. https://doi.org/10.1091/mbc.E16-03-0205.

Hosokawa, N., Hara, T., Kaizuka, T., Kishi, C., Takamura, A., Miura, Y., Iemura, S., Natsume, T., Takehana, K., Yamada, N., et al., 2009. Nutrient-dependent mTORC1 association with the ULK1-Atg13-FIP200 complex required for autophagy. Mol Biol Cell 20, 1981–1991. https://doi.org/10.1091/mbc.E08-12-1248.

Hwang, I., Oh, H., Santo, E., Kim, D.Y., Chen, J.W., Bronson, R.T., Locasale, J.W., Na, Y., Lee, J., Reed, S., et al., 2018. FOXO protects against age-progressive axonal degeneration. Aging Cell 17. https://doi.org/10.1111/acel.12701.

Hwang, R.Y., Zhong, L., Xu, Y., Johnson, T., Zhang, F., Deisseroth, K., and Tracey, W.D., 2007. Nociceptive neurons protect Drosophila larvae from parasitoid wasps. Curr Biol 17, 2105–2116. https://doi.org/10.1016/j.cub.2007.11.029.

Ikeya, T., Galic, M., Belawat, P., Nairz, K., and Hafen, E., 2002. Nutrient-dependent expression of insulin-like peptides from neuroendocrine cells in the CNS contributes to growth regulation in Drosophila. Curr Biol 12, 1293–1300.

Jaworski, J., Spangler, S., Seeburg, D.P., Hoogenraad, C.C., and Sheng, M., 2005. Control of dendritic arborization by the phosphoinositide-3’-kinase-Akt-mammalian target of rapamycin pathway. J Neurosci 25, 11300–11312. https://doi.org/10.1523/JNEUROSCI.2270-05.2005.

Jiang, N., Soba, P., Parker, E., Kim, C.C., and Parrish, J.Z., 2014. The microRNA bantam regulates a developmental transition in epithelial cells that restricts sensory dendrite growth. Development 141, 2657–2668. https://doi.org/10.1242/dev.107573.

Jung, C.H., Jun, C.B., Ro, S.H., Kim, Y.M., Otto, N.M., Cao, J., Kundu, M., and Kim, D.H., 2009. ULK-Atg13-FIP200 complexes mediate mTOR signaling to the autophagy machinery. Mol Biol Cell 20, 1992–2003. https://doi.org/10.1091/mbc.E08-12-1249.

Junger, M.A., Rintelen, F., Stocker, H., Wasserman, J.D., Vegh, M., Radimerski, T., Greenberg, M.E., and Hafen, E., 2003. The Drosophila forkhead transcription factor FOXO mediates the reduction in cell number associated with reduced insulin signaling. J Biol 2, 20. https://doi.org/10.1186/1475-4924-2-20.

Kim, W., Jang, Y.G., Yang, J., and Chung, J., 2017. Spatial Activation of TORC1 Is Regulated by Hedgehog and E2F1 Signaling in the Drosophila Eye. Dev Cell 42, 363–375 e364. https://doi.org/10.1016/j.devcel.2017.07.020.

Kimura, S., Noda, T., and Yoshimori, T., 2007. Dissection of the autophagosome maturation process by a novel reporter protein, tandem fluorescent-tagged LC3. Autophagy 3, 452–460. https://doi.org/10.4161/auto.4451.

Kumar, V., Zhang, M.X., Swank, M.W., Kunz, J., and Wu, G.Y., 2005. Regulation of dendritic morphogenesis by Ras-PI3K-Akt-mTOR and Ras-MAPK signaling pathways. J Neurosci 25, 11288–11299. https://doi.org/10.1523/JNEUROSCI.2284-05.2005.

Kwon, M.J., Han, M.H., Bagley, J.A., Hyeon, D.Y., Ko, B.S., Lee, Y.M., Cha, I.J., Kim, S.Y., Kim, D.Y., Kim, H.M., et al., 2018. Coiled-coil structure-dependent interactions between polyQ proteins and Foxo lead to dendrite pathology and behavioral defects. Proc Natl Acad Sci U S A 115, E10748–E10757. https://doi.org/10.1073/pnas.1807206115.

Lee, J.H., Cho, U.S., and Karin, M., 2016. Sestrin regulation of TORC1: Is Sestrin a leucine sensor? Sci Signal 9, re5. https://doi.org/10.1126/scisignal.aaf2885.

Martina, J.A., Chen, Y., Gucek, M., and Puertollano, R., 2012. MTORC1 functions as a transcriptional regulator of autophagy by preventing nuclear transport of TFEB. Autophagy 8, 903–914. https://doi.org/10.4161/auto.19653.

Nagel, J., Delandre, C., Zhang, Y., Forstner, F., Moore, A.W., and Tavosanis, G., 2012. Fascin controls neuronal class-specific dendrite arbor morphology. Development 139, 2999–3009. https://doi.org/10.1242/dev.077800.

Nezis, I.P., Shravage, B.V., Sagona, A.P., Lamark, T., Bjorkoy, G., Johansen, T., Rusten, T.E., Brech, A., Baehrecke, E.H., and Stenmark, H., 2010. Autophagic degradation of dBruce controls DNA fragmentation in nurse cells during late Drosophila melanogaster oogenesis. J Cell Biol 190, 523–531. https://doi.org/10.1083/jcb.201002035.

Oldham, S., Stocker, H., Laffargue, M., Wittwer, F., Wymann, M., and Hafen, E., 2002. The Drosophila insulin/IGF receptor controls growth and size by modulating PtdInsP(3) levels. Development 129, 4103–4109.

Paik, J.H., Ding, Z., Narurkar, R., Ramkissoon, S., Muller, F., Kamoun, W.S., Chae, S.S., Zheng, H., Ying, H., Mahoney, J., et al., 2009. FoxOs cooperatively regulate diverse pathways governing neural stem cell homeostasis. Cell Stem Cell 5, 540–553. https://doi.org/10.1016/j.stem.2009.09.013.

Parrish, J.Z., Xu, P., Kim, C.C., Jan, L.Y., and Jan, Y.N., 2009. The microRNA bantam functions in epithelial cells to regulate scaling growth of dendrite arbors in drosophila sensory neurons. Neuron 63, 788–802. https://doi.org/10.1016/j.neuron.2009.08.006.

Poe, A.R., Tang, L., Wang, B., Li, Y., Sapar, M.L., and Han, C., 2017. Dendritic space-filling requires a neuronal type-specific extracellular permissive signal in Drosophila. Proc Natl Acad Sci U S A 114, E8062–E8071. https://doi.org/10.1073/pnas.1707467114.

Poe, A.R., Wang, B., Sapar, M.L., Ji, H., Li, K., Onabajo, T., Fazliyeva, R., Gibbs, M., Qiu, Y., Hu, Y., et al., 2019. Robust CRISPR/Cas9-Mediated Tissue-Specific Mutagenesis Reveals Gene Redundancy and Perdurance in Drosophila. Genetics 211, 459–472. https://doi.org/10.1534/genetics.118.301736.

Puig, O., Marr, M.T., Ruhf, M.L., and Tjian, R., 2003. Control of cell number by Drosophila FOXO: downstream and feedback regulation of the insulin receptor pathway. Genes Dev 17, 2006–2020. https://doi.org/10.1101/gad.1098703.

Rajan, A., and Perrimon, N., 2012. Drosophila cytokine unpaired 2 regulates physiological homeostasis by remotely controlling insulin secretion. Cell 151, 123–137. https://doi.org/10.1016/j.cell.2012.08.019.

Renault, V.M., Rafalski, V.A., Morgan, A.A., Salih, D.A., Brett, J.O., Webb, A.E., Villeda, S.A., Thekkat, P.U., Guillerey, C., Denko, N.C., et al., 2009. FoxO3 regulates neural stem cell homeostasis. Cell Stem Cell 5, 527–539. https://doi.org/10.1016/j.stem.2009.09.014.

Roczniak-Ferguson, A., Petit, C.S., Froehlich, F., Qian, S., Ky, J., Angarola, B., Walther, T.C., and Ferguson, S.M., 2012. The transcription factor TFEB links mTORC1 signaling to transcriptional control of lysosome homeostasis. Sci Signal 5, ra42. https://doi.org/10.1126/scisignal.2002790.

Rulifson, E.J., Kim, S.K., and Nusse, R., 2002. Ablation of insulin-producing neurons in flies: growth and diabetic phenotypes. Science 296, 1118–1120. https://doi.org/10.1126/science.1070058.

Ruvinsky, I., and Meyuhas, O., 2006. Ribosomal protein S6 phosphorylation: from protein synthesis to cell size. Trends Biochem Sci 31, 342–348. https://doi.org/10.1016/j.tibs.2006.04.003.

Sancak, Y., Thoreen, C.C., Peterson, T.R., Lindquist, R.A., Kang, S.A., Spooner, E., Carr, S.A., and Sabatini, D.M., 2007. PRAS40 is an insulin-regulated inhibitor of the mTORC1 protein kinase. Mol Cell 25, 903–915. https://doi.org/10.1016/j.molcel.2007.03.003.

Schaffner, I., Minakaki, G., Khan, M.A., Balta, E.A., Schlotzer-Schrehardt, U., Schwarz, T.J., Beckervordersandforth, R., Winner, B., Webb, A.E., DePinho, R.A., et al., 2018. FoxO Function Is Essential for Maintenance of Autophagic Flux and Neuronal Morphogenesis in Adult Neurogenesis. Neuron 99, 1188–1203 e1186. https://doi.org/10.1016/j.neuron.2018.08.017.

Sears, J.C., and Broihier, H.T., 2016. FoxO regulates microtubule dynamics and polarity to promote dendrite branching in Drosophila sensory neurons. Dev Biol 418, 40–54. https://doi.org/10.1016/j.ydbio.2016.08.018.

Settembre, C., Di Malta, C., Polito, V.A., Garcia Arencibia, M., Vetrini, F., Erdin, S., Erdin, S.U., Huynh, T., Medina, D., Colella, P., et al., 2011. TFEB links autophagy to lysosomal biogenesis. Science 332, 1429–1433. https://doi.org/10.1126/science.1204592.

Skalecka, A., Liszewska, E., Bilinski, R., Gkogkas, C., Khoutorsky, A., Malik, A.R., Sonenberg, N., and Jaworski, J., 2016. mTOR kinase is needed for the development and stabilization of dendritic arbors in newly born olfactory bulb neurons. Dev Neurobiol 76, 1308–1327. https://doi.org/10.1002/dneu.22392.

Tang, H.Y., Smith-Caldas, M.S., Driscoll, M.V., Salhadar, S., and Shingleton, A.W., 2011. FOXO regulates organ-specific phenotypic plasticity in Drosophila. PLoS Genet 7, e1002373. https://doi.org/10.1371/journal.pgen.1002373.

Tracey, W.D., Jr., Wilson, R.I., Laurent, G., and Benzer, S., 2003. painless, a Drosophila gene essential for nociception. Cell 113, 261–273. https://doi.org/10.1016/s0092-8674(03)00272-1.

Vaadia, R.D., Li, W., Voleti, V., Singhania, A., Hillman, E.M.C., and Grueber, W.B., 2019. Characterization of Proprioceptive System Dynamics in Behaving Drosophila Larvae Using High-Speed Volumetric Microscopy. Curr Biol 29, 935–944 e934. https://doi.org/10.1016/j.cub.2019.01.060.

Vander Haar, E., Lee, S.I., Bandhakavi, S., Griffin, T.J., and Kim, D.H., 2007. Insulin signalling to mTOR mediated by the Akt/PKB substrate PRAS40. Nat Cell Biol 9, 316–323. https://doi.org/10.1038/ncb1547.

Verdu, J., Buratovich, M.A., Wilder, E.L., and Birnbaum, M.J., 1999. Cell-autonomous regulation of cell and organ growth in Drosophila by Akt/PKB. Nat Cell Biol 1, 500–506. https://doi.org/10.1038/70293.

Wang, L., Harris, T.E., Roth, R.A., and Lawrence, J.C., Jr., 2007. PRAS40 regulates mTORC1 kinase activity by functioning as a direct inhibitor of substrate binding. J Biol Chem 282, 20036–20044. https://doi.org/10.1074/jbc.M702376200.

Watanabe, K., Furumizo, Y., Usui, T., Hattori, Y., and Uemura, T., 2017. Nutrient-dependent increased dendritic arborization of somatosensory neurons. Genes Cells 22, 105–114. https://doi.org/10.1111/gtc.12451.

Xiang, Y., Yuan, Q., Vogt, N., Looger, L.L., Jan, L.Y., and Jan, Y.N., 2010. Light-avoidance-mediating photoreceptors tile the Drosophila larval body wall. Nature 468, 921–926. https://doi.org/10.1038/nature09576.

Yeo, H., Lyssiotis, C.A., Zhang, Y., Ying, H., Asara, J.M., Cantley, L.C., and Paik, J.H., 2013. FoxO3 coordinates metabolic pathways to maintain redox balance in neural stem cells. EMBO J 32, 2589–2602. https://doi.org/10.1038/emboj.2013.186.

Zhang, T., Zhou, Q., Ogmundsdottir, M.H., Moller, K., Siddaway, R., Larue, L., Hsing, M., Kong, S.W., Goding, C.R., Palsson, A., et al., 2015. Mitf is a master regulator of the v-ATPase, forming a control module for cellular homeostasis with v-ATPase and TORC1. J Cell Sci 128, 2938–2950. https://doi.org/10.1242/jcs.173807.

Zoncu, R., Efeyan, A., and Sabatini, D.M., 2011. mTOR: from growth signal integration to cancer, diabetes and ageing. Nat Rev Mol Cell Biol 12, 21–35. https://doi.org/10.1038/nrm3025.

